# Impact of pregravid obesity on anti-microbial fetal monocyte response

**DOI:** 10.1101/2022.07.10.499492

**Authors:** Suhas Sureshchandra, Brianna M. Doratt, Norma Mendoza, Oleg Varlamov, Monica Rincon, Nicole E. Marshall, Ilhem Messaoudi

**Author notes:** Corresponding Author: Ilhem Messaoudi Microbiology, Immunology, and Molecular Genetics College of Medicine University of Kentucky.

## Abstract

Maternal pre-pregnancy (pregravid) obesity is associated with adverse outcomes for both mother and offspring. Amongst the complications for the offspring is increased susceptibility and severity of neonatal infections necessitating admission to the intensive care unit, notably bacterial sepsis and enterocolitis. Previous studies have reported aberrant responses to LPS and polyclonal stimulation by umbilical cord blood monocytes that were mediated by alterations in the epigenome. In this study, we show that pregravid obesity dysregulates umbilical cord blood monocyte responses to bacterial and viral pathogens. Specifically, interferon-stimulated gene expression and inflammatory responses to *E. coli* and respiratory syncytial virus (RSV) were significantly dampened. Although upstream signaling events were comparable, translocation of the key transcription factor NF-κB and chromatin accessibility at pro- inflammatory gene promoters following TLR stimulation was significantly attenuated. Using a rhesus macaque model of western style diet induced obesity, we further demonstrate that this defect is detected in fetal peripheral monocytes and tissue-resident macrophages during gestation. Collectively, these data indicate that maternal obesity and high-fat diet present metabolic, signaling, and epigenetic impediments to pathogen recognition in fetal innate immune cells that result in a state of immune paralysis during gestation and at birth.

## INTRODUCTION

Nearly 40% of women of childbearing age (18-44 years old) in the United States met body mass index (BMI) obesity criteria in 2019 (*1*) and the rate of obesity in women of childbearing age continues to rise with ∼11% increase in prevalence from 2016-2019 (*2*). Consequently, pre- pregnancy (pregravid) obesity has emerged as one of the leading co-morbidities that impact maternal and fetal health. The Developmental Origins of Health and Disease (DOHaD) concept (*3, 4*) proposes that both maternal overnutrition and undernutrition during fetal development have a programming effect on offspring’s subsequent responses to the environment during childhood and adult life (*5, 6*). Indeed, it is now well established that pregravid obesity is associated with adverse outcomes for both mother and fetus with long-term health consequences for the offspring (*7, 8*). Compared to lean mothers, mothers with obesity have a substantially increased risk of developing gestational diabetes, pre-eclampsia, and hypertension all associated with small for gestational age neonates and increased risk of stillbirth (*9, 10*). Pregravid obesity is linked to a higher incidence of long-term consequences for the offspring including increased incidence of allergies, asthma, and metabolic disorders (*11–21*). Additionally, maternal obesity is associated with an increased risk of necrotizing enterocolitis, sepsis, and severe respiratory syncytial virus (RSV) infection necessitating admission to the neonatal intensive care unit (*22–24*).

Similarly, studies using animal models where maternal obesity is induced via the administration of high fat or western style diets (HFD, WSD) showed that pups born to obese dams fed a HFD during gestation generated lower ovalbumin (OVA)-specific IgG, but higher OVA-specific IgE, after immunization with OVA compared to pups born to dams fed a control diet (*25*). These data potentially explain the increased incidence of asthma/wheezing in children born to mothers with obesity (*13*). Further studies show that pups born to obese dams fed a HFD are more susceptible to bacterial challenge (*26*), develop more severe bronchiolitis following RSV challenge (*22*), and are more likely to develop autoimmune encephalitis (*26*). Moreover, ex vivo LPS stimulation of colonic lamina propria lymphocytes (LPLs) isolated from pups born to dams fed a HFD resulted in greater levels of IL-6, IL-1β, and IL-17, whereas LPS stimulation of splenocytes from the same animals generated decreased levels of TNFα and IL-6 compared to control groups (*26*).

These observations suggest that pregravid obesity disrupts the development and maturation of the offspring’s immune system *in utero.* Indeed, clinical studies showed that high pregravid BMI is associated with reduced frequency of umbilical cord blood (UCB) CD4^+^ T cells and dampened CD4^+^ T cell responses to polyclonal stimulation (*15, 27–29*). Additionally, pregravid obesity was associated with a reduced ability of human UCB CD14^+^ monocytes to respond to ex vivo stimulation with toll-like receptor (TLR) ligands (*15, 28*). Moreover, monocytes from offspring of mothers with obesity showed a decrease in transcripts of the pro-inflammatory cytokines IL-1ß and IL-12ß (*30*). These dampened CD4^+^ T cell and monocyte responses are mediated by epigenetic changes at key loci that regulate inflammatory responses and cellular differentiation pathways (*27, 28*). Other studies have reported an increased expression of PPAR-γ, which can attenuate macrophage inflammatory responses (*31*) in peripheral blood mononuclear cells (PBMCs) obtained from children born to mothers with obesity (*32*). Similarly, we reported hypomethylation of the PPAR-γ gene in UCB monocytes obtained from mothers with obesity, thus indicating sustained reprogramming (*28*). Alterations in the DNA methylation patterns of the promoter regions in monocytes of the obese group suggest dysregulated responses to M1 and M2 macrophage polarization stimuli (*28*).

In this study, we extended these earlier clinical studies and interrogated the impact of pregravid obesity on the response to bacterial and viral pathogens. Our data showed dysregulated response by UCB monocytes from babies born to mothers with obesity. Additional analyses showed that upstream signaling events were comparable between UCB from lean and obese mothers, however, reduced chromatin accessibility and lack of chromatin remodeling in response to stimulation resulted in a dampened transcriptional response. This dampened response was also detected in PBMC obtained from neonatal rhesus macaque born to obese dams fed a high fat diet indicating that these observations are not unique to UCB.

## METHODS

### Subjects and experimental design

This study was approved by the Institutional Ethics Review Board of Oregon Health and Science University and the University of California, Irvine. Written consent was obtained from all subjects. The mean age of the 43 lean women was 33.1 ± 4.4 years with an average pre-pregnancy BMI of 21.9 ± 1.7 kg/m^2^ and gestation age at delivery of 39.6 ± 1.4 weeks; mean age of the 36 obese women was 30.7 ± 4.9 years with an average pre-pregnancy BMI of 38.2 ± 8.4 kg/m^2^ and gestation age at delivery of 39.3 ± 1.4 weeks (Table 1). The racial distribution of the lean cohort was 2.32% Asian American, 6.97% Hispanic, 88.37% Caucasian, and 2.32% declined to report. The obese cohort consisted of 25.00% Hispanic, 66.67% Caucasian, 2.77% with more than one race, and 5.55% who declined to report. Exclusion criteria included active maternal infection, documented fetal congenital anomalies, substance abuse, chronic illness requiring regular medication use, preeclampsia, gestational diabetes, chorioamnionitis, and significant medical conditions (active cancers, cardiac, renal, hepatic, or pulmonary diseases), or an abnormal glucose tolerance test. Women underwent a fasting blood draw and body composition via air displacement plethysmography using a BodPod (Life Measurement Inc, Concord, CA).

**Table 1:**
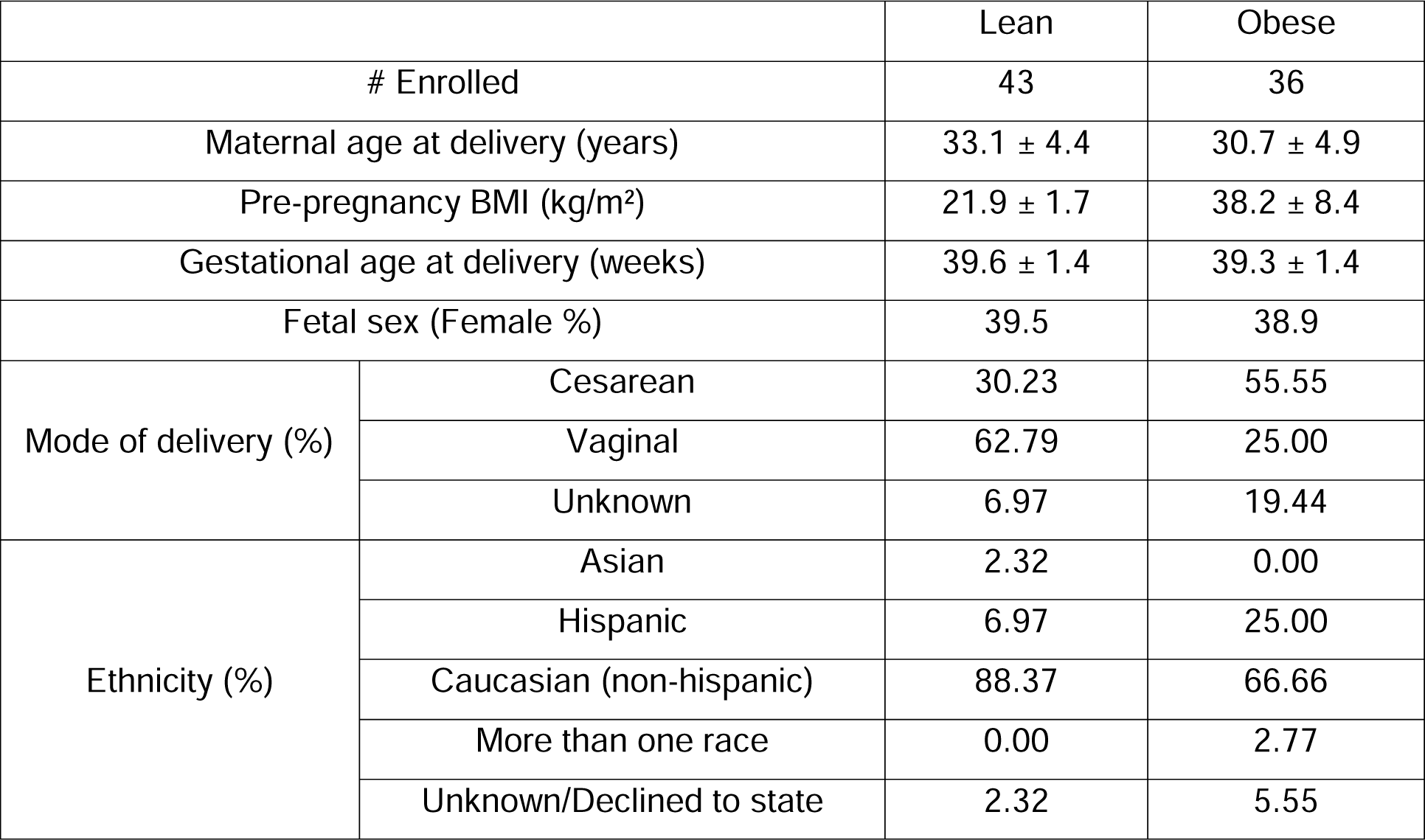
Cohort characteristics

### Umbilical Cord Blood Mononuclear Cell (UCBMC) isolation

Complete blood counts were obtained by Beckman Coulter Hematology analyzer (Brea, CA) before cell isolation. Umbilical cord blood mononuclear cells (UCBMC) and plasma were obtained by standard density gradient centrifugation over Ficoll (GE Healthcare, Chicago, IL). UCBMC were frozen in 10% DMSO/FetalPlex (GeminiBio, Sacramento, CA) and stored in liquid nitrogen until analysis. Plasma was stored at -80°C until analysis.

### Cord Blood Immunophenotyping

10^6^ UCBMC were stained using the following cocktail of antibodies to enumerate innate immune cells and their subsets: PE-CD3, PE-CD19, PB-CD16, PE-Cy7-CD11c, AF700-CD14, PCP- Cy5.5-CD123, BV711-CD56, and APC-Cy7-HLA-DR. All samples were acquired with the Attune NxT Flow Cytometer (ThermoFisher Scientific, Waltham, MA) and analyzed using FlowJo 10.5.0 (Ashland, OR).

### Intracellular cytokine assay

To measure cytokine responses of monocytes and dendritic cells, 10^6^ UCBMC were stimulated for 16h at 37°C in RPMI 1640 medium supplemented with 10% FBS in the presence or absence of 1 ug/mL LPS (TLR4 ligand, *E. Coli* 055:B5; Invivogen, San Diego CA). Brefeldin A (Sigma, St. Louis MO) was added after 1-hour incubation. Cells were stained for APC-Cy7-CD14 and PCP-Cy5.5-HLA-DR, fixed, permeabilized, and stained intracellularly for APC-TNFα and PE-IL-6. All samples were acquired with the Attune NxT Flow Cytometer (ThermoFisher Scientific, Waltham, MA) and analyzed using FlowJo 10.5.0 (Ashland, OR).

### Pathogen stimulation

Approximately 5x10^5^ MACS purified UCB monocytes were cultured with RSV (Human respiratory syncytial virus ATCC® VR-1540, Manassas, VA) or *E. Coli* (Escherichia coli (Migula) Castellani and Chalmers ATCC 11775, Manassas, VA) or left untreated in RP10 medium for 16 hours at 37°C. RSV was added at a multiplicity of infection (MOI) of 5 and *E. coli* at 6x10^5^ cfu/mL. Following the 16hr incubation, cells were spun down. Cell pellets were frozen in QIAzol lysis reagent (Qiagen, Hilden Germany) to generate RNA-Seq libraries. Cell supernatants were frozen at -80C to measure the concentrations of chemokines and cytokines using Luminex.

### Luminex and ELISA

Immune mediators in plasma were measured using a customized multiplex human factor panel (R & D Systems, Minneapolis MN) measuring cytokines (IFNβ, IFNγ, IL-1β, IL-10, IL-12p70, IL- 13, IL-15, IL-17A, IL-18, IL-1RA, IL-2, IL-21, IL-4, IL-5, IL-7, TNFα, IL-23, IL-31, IL-22, IL-27), chemokines (CCL2/MCP-1, CCL3/MIP-1α, CCL4/MIP-1β, CCL5/RANTES, CCL11/Eotaxin, CXCL1/GROα, CXCL8/IL-8, CXCL9/MIG, CXCL10/IP-10, CXCL11/I-TAC, CXCL12/SDF-1α, CXCL13/BCA-1), growth factors (BDNF, GM-CSF, HGF, EGF, VEGF, PDGF-BB) and additional molecules (PD-L1, S100). Metabolic hormones were measured using a 3-plex kit measuring insulin, leptin, and PYY (Millipore, Burlington MA). Adipokines were assayed using a 5-plex kit measuring adiponectin, adipsin, lipocalin-2, total PAI-1, and resistin (Millipore, Burlington MA). CRP and IL-6 were measured in UCB plasma using a high-sensitivity ELISA (Life Technologies, Carlsbad CA) per the manufacturer’s instructions.

Supernatants from fetal rhesus macaque monocyte stimulation experiments were analyzed using an NHP XL Cytokine Premixed 36-plex kit (Bio-Techne, Minneapolis MN). Samples were diluted per the manufacturer’s instructions and analyzed in duplicate on the Magpix Instrument (Luminex, Austin, TX). Data were fit using a 5P-logistic regression on xPONENT software.

### Bulk RNA-Seq

Total RNA was isolated from monocytes using an mRNeasy kit (Qiagen, Valencia CA). Quality and concentrations were measured using Agilent 2100 Bioanalyzer. Libraries were generated using the TruSeq Stranded Total RNA-Seq kit (Illumina, San Diego CA). Briefly, following rRNA depletion, mRNA was fragmented for 8 min, converted to double-stranded cDNA, and adapter ligated. Fragments were then enriched by PCR amplification and purified. The size and quality of the library were verified using Qubit and Bioanalyzer. Libraries were multiplexed and sequenced on the HiSeq4000 platform (Illumina, San Diego CA) to yield an average of 20 million 100 bp single-end reads per sample.

### Bulk RNA-Seq analysis

Quality control of raw reads was performed using FASTQC retaining bases with quality scores of ≥20 and reads ≥35 base pairs long. Reads were aligned to the human genome (hg38) using splice-aware aligner TopHat using annotations available from ENSEMBL (GRCh38.85) database. Lowly expressed genes were filtered at the counting stage, eliminating genes with 0 counts in at least 3 samples regardless of the group. Quantification of read counts was performed using the GenomicRanges package in R and normalized to derive transcripts per million (TPM) counts.

Responses to RSV and *E. coli* were modeled pairwise relative to unstimulated samples using negative binomial GLMs following low read count filtering. Genes with log_2_ fold change ≥ 1 in either direction and corrected expression difference FDR<0.05 were considered differentially expressed genes (DEG). Functional enrichment of DEG was performed using Metascape (*33*). Heatmaps of fold change or TPMs and bubble plots of enrichment of Gene Ontology (GO) terms were generated using ggplot in R.

### Cell Sorting and library generation for single cell (sc)RNA-seq

UCBMC were thawed then washed twice in PBS with 0.04% BSA and incubated with individual 3’ CellPlex oligos (10X Genomics) per manufacturer’s instructions for 5 minutes at room temperature. Pellets were washed three times in PBS with 1% BSA, resuspended in 300 uL FACS buffer, and sorted on BD FACS Aria Fusion into RPMI (supplemented with 30% FBS) following the addition of SYTOX Blue stain (1:1000, ThermoFisher) for live versus dead discrimination. Sorted cells were counted in triplicates and resuspended in PBS with 0.04% BSA in a final concentration of 1200 cells/uL. Cells were immediately loaded in the 10X Genomics Chromium with a loading target of 17,600 cells. Libraries were generated using the V3.1 chemistry (gene expression) and Single Cell 3L Feature Barcode Library Kit per the manufacturer’s instructions (10X Genomics, Pleasanton CA). Libraries were sequenced on Illumina NovaSeq 6000 with a sequencing target of 50,000 gene expression reads and 5,000 multiplexed CellPlex reads per cell.

### scRNA-seq data analysis

For 3’ gene expression with CellPlex, raw reads were aligned and quantified using Cell Ranger (version 6.0.2, 10X Genomics) against the human reference genome (GRCh38) using the *multi* option and CMO information. Only singlets identified from each sample were included in downstream analyses. Droplets with ambient RNA (cells fewer than 400 detected genes), potential doublets (cells with more than 4000 detected genes) and dying cells (cells with more than 20% total mitochondrial gene expression) were excluded during initial QC. Data objects from lean and obese groups were integrated using Seurat. Data normalization and variance stabilization was performed using the *SCTransform* function using a regularized negative binomial regression, correcting for differential effects of mitochondrial and ribosomal gene expression levels and cell cycle. Dimension reduction was performed using the *RunPCA* function to obtain the first 30 principal components followed by clustering using the *FindClusters* function in Seurat. Visualization of clusters was performed using the UMAP algorithm as implemented by Seurat’s *runUMAP* function. Cell types were assigned to individual clusters using the *FindMarkers* function with a fold change cutoff of at least 0.5 and using a known catalog of well-characterized scRNA markers for PBMC (*34*).

Differential expression analysis was tested using the Wilcoxon rank sum test followed by Bonferroni correction using all genes in the dataset. For gene scoring analysis, we compared gene signatures and pathways from KEGG (https://www.genome.jp/kegg/pathway.html) in subpopulations using Seurat’s *AddModuleScore* function. All graphs were generated in R.

### Phagocytosis Assay

To quantify the phagocytic ability of UCB monocytes, monocytes were isolated using magnetically activated cell sorting and anti-CD14 antibody coupled to magnetic beads per the manufacturer’s recommendation (MACS, Miltenyi Biotech, San Jose, CA). Cells were activated with 1ug/ml LPS for 16 hours, then cultured with *E. coli* particles conjugated with HRP (horseradish peroxidase) in 96-well plates per manufacturer’s protocol (CytoSelect 96-well phagocytosis assay, Cell Biolabs, San Diego CA) for 3 hours in 37°C incubator with 5% CO2.

Cells were washed, fixed, permeabilized, incubated with substrate, and quantified using colorimetry (CytoSelect 96-well phagocytosis assay, Cell Biolabs, San Diego CA).

### Cell Migration Assay

The migratory potential of monocytes to supernatants containing chemokines was measured using the CytoSelect 96-well Cell Migration Assay Cell Migration assay (Cell Biolabs, San Diego CA). Briefly, 2x10^5^ purified UCB monocytes isolated using anti-CD14 antibody coupled to magnetic beads per the manufacturer’s recommendation (MACS, Miltenyi Biotech, San Jose, CA) were incubated in serum-free media in the upper wells of the migration plate, while supernatants collected following PMA (phorbol myristate acetate) stimulation of adult PBMC were placed in lower wells and incubated at 37°C and 5% CO_2_ for 5 hr. The number of cells that migrated into the lower wells was quantified using CyQuant cell proliferation assay per the manufacturer’s instructions. Absolute numbers of migrated cells were calculated using a standard curve for CyQuant assay with a linear range of fluorescence limited from 50 to 50,000 cells. Cell-free media served as a negative control.

### Bulk ATAC-Seq

ATAC-Seq libraries were generated using OMNI-ATAC to reduce mitochondrial reads (*35*). Briefly, 50,000 purified UCB monocytes were stimulated with 1 ug/mL LPS for 16 hours before being lysed in lysis buffer (10mM Tris-HCl (pH 7.4), 10 mM NaCl, 3 mM MgCl_2_), for 3 min on ice to prepare the nuclei. Immediately after lysis, nuclei were spun at 500g for 10 min to remove the supernatant. Nuclei were then incubated with a transposition mixture (100 nM Tn5 transposase, 0.1% Tween-20, 0.01% Digitonin, and TD Buffer) at 37C for 30 min. Transposed DNA was then purified with AMPure XP beads (Beckman Coulter) and partially amplified for 5 cycles using the following PCR conditions - 72°C for 3 min; 98°C for 30s and thermocycling at 98°C for 10s, 63°C for 30s, and 72°C for 1 min. To avoid overamplification, qPCR was performed on 5 uL of partially amplified library. Additional cycles of amplification for the remainder of the sample were calculated from the saturation curves (cycles corresponding to a third of the saturation value). Fully amplified samples were purified with AMPure beads and quantified on the Bioanalyzer (Agilent Technologies, Santa Clara CA). Libraries were sequenced on the HiSeq4000 platform (Illumina, San Diego CA).

### Analysis of bulk ATAC-Seq data

Paired reads from sequencing were quality-checked using FASTQC and trimmed to retain reads with quality scores of ≥20 and minimum read lengths of 50 bp. Trimmed paired reads were aligned to the human genome (hg38) using Bowtie2 (–X 2000 –k 1 –very-sensitive –no- discordant –no-mixed). Reads aligning to mitochondrial genome and allosomes were removed using samtools. PCR duplicate artifacts were then removed using Picard. Finally, poor quality alignments and improperly mapped reads were filtered using samtools (samtools view –q 10 –F 3844). To reflect the accurate read start site due to Tn5 insertion, BAM files were repositioned using the ATACseqQC package in R. The positive and negative strands were offset by +4bp and -5bp respectively. Samples within a group were merged and sorted using samtools.

Sample QC and statistics for merged BAM files were generated using HOMER makeTagDirectory function. Accessible chromatin peaks were called for mapped paired reads using the HOMER findpeak function (-minDist 150 –region –fdr 0.05). Differentially accessible regions (DAR) in either direction were captured using HOMER getDiffererentialPeaks function (- q 0.05). DAR were annotated using the human GTF annotation file (GRCh38.85) and ChIPSeeker with a promoter definition of -1000 bp and +100 bp around the transcriptional start site (TSS). Peaks overlapping 5’UTRs, promoters, first exons, and first introns were pooled for functional enrichment of genes. For intergenic changes, the genes closest to the intergenic DAR were considered. Functional enrichment of this pooled list of genes was performed using DAVID (Fisher p-value <0.05). BAM files were converted to bigWig files using bedtools and visualized on the new WashU EpiGenome browser.

### scATAC sample preparation

One to two million UCBMC (n=4/group) were incubated for 4 hours at 37°C in the presence or absence of 1 ug/mL LPS. Cell pellets were then washed, and surface-stained with monocyte markers CD14-AF700 and HLA-DR APC-Cy7 for 30 minutes. Stained cells were washed, resuspended in FACS buffer, and stained for live-dead exclusion (SYTOX Blue, 1:1000 dilution). Equal numbers of monocytes (CD14+HLA-DR+) were sorted and pooled by group (lean/obese) into RPMI supplemented with 30% FBS. Cells were washed thoroughly, and nuclei were isolated using the low cell input nuclei isolation protocol (10X Genomics). Nuclei were counted and verified for integrity then resuspended in PBS with 0.04% BSA at 1000 nuclei/uL concentration. Monocyte nuclei were loaded onto the 10X Genomics Chromium according to the manufacturer’s protocol using the single-cell ATAC kit (v2). Library preparation was performed per the manufacturer’s protocol prior to sequencing on Illumina NovaSeq 6000 platform.

### scATAC data analysis

Basecall files were used to generate FASTQ files with cellranger-atac (v1; 10X Genomics). Reads were aligned to the human genome using cellranger-atac count with the cellranger-atac- GRCh38.1.1.0 reference. Mapped Tn5 insertion sites from cellranger were read into ArchR (version 1.0.1) R package retaining barcodes with at least 1000 fragments per cell and a TSS enrichment score > 4. Doublets were identified and filtered using *addDoubletScores* and *filterDoublets* (filter ratio = 1.4) respectively before iterative LSI dimensionality reduction (iterations = 2, res = 0.2, variable features = 25000, dim = 30). Clustering was then performed (*addClusters*, res = 0.8) before UMAP dimensionality reduction (nNeighbors = 30, metric = cosine, minDist = 0.4). One cluster enriched for high doublet scores and was removed. Peaks for each cluster were calculated using MACS2, using the *addReproduciblePeakSet* function. Marker peaks for each cluster and differential peaks with stimulation were calculated using the *getMarkerFeatures* function using the Wilcoxon test. A cluster of activated monocytes was identified by pileups and feature plots of canonical cytokine and activation markers.

### Metabolic Assays

Oxygen Consumption Rate (OCR) and Extracellular Acidification Rate (ECAR) were measured using Seahorse XF Glycolysis Rate Assay on Seahorse XFp Flux Analyzer (Agilent Technologies, Santa Clara, CA) following the manufacturer’s instructions. Briefly, 200,000 purified monocytes (pooled n=3/group) were seeded in glucose-free media and cultured in the presence/absence of 1 ug/mL LPS for 1 hour in a 37°C incubator without CO_2_ on Cell-Tak (Corning, Corning, NY) coated 8-well culture plates in phenol-free RPMI media containing 2 mM L-glutamine, 1mM sodium pyruvate, and 5mM HEPES. Plates were run on the XFp for 8 cycles of basal measurements, followed by acute injection of L-glucose (100 mM), oligomycin (50 uM), and 2-DG (500 mM). Data were analyzed on Seahorse Wave desktop software (Agilent Technologies, Santa Clara, CA).

### Histone ELISA

Nuclear extracts from 2L×L10^5^ UCB monocytes purified using an anti-CD14 antibody coupled to magnetic beads per the manufacturer’s recommendation (MACS, Miltenyi Biotech, San Jose, CA) were isolated per the manufacturer’s instructions (Abcam, Cambridge UK) and quantified using a micro-BCA assay protein assay kit (ThermoFisher Scientific, Waltham, MA). Histone modifications were measured using a Histone H3 modification Multiplex Assay (Abcam, Cambridge UK). The input was normalized based on total protein concentration, and 20Lng of nuclear extract was added to each well. Given limited sample availability, only a subset of histone methylation marks were assayed (H3K4me1, H3K4me2, H3K4me3, H3K9me3, H3K9Ac, and total H3). Optical density was measured at 450 nm. All values are reported as percentages of the total H3 signal.

### Histone Flow

Activation-induced changes in histone post-translational modifications were assayed using flow cytometry (n=8/group). Briefly, 10^6^ UCBMC were stimulated with 1 μg/mL LPS for 2 hours in a 37°C incubator with 5% CO2. Cell pellets were washed with FACS buffer, surface stained for monocytes (CD14 AF700, HLA-DR APC-Cy7) for 20 minutes, washed, and fixed using Foxp3/Transcription factor Fix/Perm buffer (Tonbo Biosciences, San Diego, CA) for 1h at 4°C. Pellets were then stained intracellularly overnight with antibodies against H3K4me3-AF647 (Clone EPR20551-225, Abcam, Cambridge UK) and H3K9me3-PE (Clone D4W1U, Cell Signaling Technology, Danvers, MA). Cells were washed twice with Perm buffer (Tonbo Biosciences, San Diego, CA), followed by FACS buffer. After the final wash, pellets were resuspended in FACS buffer and analyzed on Attune NxT flow cytometer (ThermoFisher Scientific, Waltham, MA).

### Phospho-Flow

Activation-induced changes in signaling mediators downstream of TLR4 were measured using flow cytometry. Briefly, 10^6^ UCBMC (n=8/group) were stimulated either for 30 minutes or 2 hours with 1 μg/mL LPS in a 37°C incubator with 5% CO2. Cells were washed with FACS buffer and surface stained for CD14 and HLA-DR in FACS tubes. Pellets were washed in FACS buffer and resuspended in 100 μL prewarmed PBS (Ca+ Mg+ free). Cells were fixed immediately by the addition of equal volumes of prewarmed Cytofix Buffer (BD Biosciences, Brea, CA) and thorough mixing and incubating at 37°C for 10 minutes. Cells were then centrifuged at 600g for 8 minutes. Supernatants were removed leaving no more than 50 μL residual volume. Cells were then permeabilized by the addition of 1 mL 1X BD PermWash Buffer I, mixed well, and incubated at room temperature for 30 minutes. Pellets were then washed and stained intranuclearly with antibodies against NF-kB p65 pS529 AF647 (Clone K10-895.12.50, Cell Signaling Technology, Danvers, MA) (for 30 minutes stimulation samples) or IkBa PE (Clone MFRDTRK, eBioscience, San Diego CA) and Phospho-p38 MAPK-APC (Clone 4NIT4KK, eBioscience, San Diego CA) for 1h at room temperature in the dark. Samples were washed twice in Permwash Buffer I, resuspended in FACS buffer, and analyzed on Attune NxT flow cytometer.

### Imaging flow cytometry

Nuclear translocation of p50 was measured using the Amnis NFkB Translocation kit (Luminex Corporation, Austin TX). Briefly, 1-2 x 10^6^ UCBMC were stimulated with 1μg/mL LPS for 4 hours in a 37°C incubator with 5% CO2. Cell pellets were washed thoroughly and stained for dead cell exclusion (Ghost Violet 510, 1:1000 dilution Tonbo Biosciences, San Diego, CA) for 30 minutes at 4°C. Cells were then washed, and surface stained for 20 minutes at 4°C (CD14-APC, HLA-DR-APC-Cy7). Cells were washed in 1X Assay Buffer (Luminex Corporation, Austin TX) and fixed using 1X Fixation buffer for 10 minutes at room temperature. Washed pellets were then stained intranuclearly for p50-AF488 (1:20 dilution) in 1X Assay Buffer for 30 minutes at room temperature in the dark. At the end of the incubation, cells were washed twice in 1X Assay buffer and resuspended in 50 μL 0.25X Fixation Buffer in polypropylene Eppendorf tubes. Samples were analyzed on the Amnis Imagestream XMark II imaging flow cytometry platform and analyzed on the IDEAS Software using the nuclear translocation wizard.

### Fetal rhesus macaque samples macrophage isolation and stimulation

PBMC, ileal lamina propria lymphocytes, and splenocytes from gestational day (GD) 130 rhesus macaque fetuses born to obese and lean dams (n=3/group) were isolated and cryopreserved from animals described in (*36*). Cells were thawed, and surface stained for CD14 and HLA-DR. PBMC were stimulated with 1 ug/mL LPS for 16 hours in a 37°C incubator with 5% CO2. The frequency of responding monocytes was determined using intracellular cytokine staining for IL-6 and TNFL following surface staining for CD14+DR+ cells. For tissue-resident macrophages, 10,000 CD14^high^HLA-DR+ splenic macrophages and 5,000 ileal CD14^high^HLA-DR^+^ macrophages were plated per condition from each sample and stimulated with *E. coli (*6x10^5^ cfu/mL) or left untreated for 16 hours at 37°C. Plates were spun and supernatants were collected for analysis of secreted cytokines and chemokines using Luminex.

### Statistical analysis

All statistical analyses were conducted in Prism 8 (GraphPad, San Diego CA). All definitive outliers in two-way and four-way comparisons were identified using ROUT analysis (Q=0.1%). Data was then tested for normality using the Shapiro-Wilk test (alpha=0.05). If data were normally distributed across all groups, differences with obesity and pregnancy were tested using ordinary one-way ANOVA with unmatched samples. Multiple comparisons were corrected using the Holm-Sidak test adjusting the family-wise significance and confidence level at 0.05. If the Gaussian assumption was not satisfied, differences were tested using the Kruskal-W allis test (alpha=0.05) followed by Dunn’s multiple hypothesis correction tests. Differences in normally distributed datasets were tested using an unpaired t-test with Welch’s correction (assuming different standard deviations). Two group comparisons that failed normality tests were carried out using the Mann-Whitney test.

## RESULTS

### Cord blood monocytes from babies born to mothers with obesity exhibit attenuated responses to TLR stimulation

To dissect the mechanisms by which pregravid obesity impacts functional responses in cord blood monocytes, we carried a multi-pronged approach (**Figure 1A**). We began by characterizing immune cell composition and phenotypes in cord blood of subjects stratified by their pre-pregnancy body mass index (BMI) – babies born to lean mothers (BMI <25) and babies born to mothers with obesity (BMI >30). As previously reported, maternal obesity did not alter the frequencies of white blood cells (WBC) or their subsets in the cord blood (*15, 27, 28, 37*) (**Supp Figure 1A**). Additionally, no changes were observed in cord blood plasma levels of inflammatory markers (IL-6, CRP) or insulin (**Supp Figure 1B-C**). Leptin levels were, however, significantly higher in the obese group (**Supp Figure 1C**).

**Figure 1:**
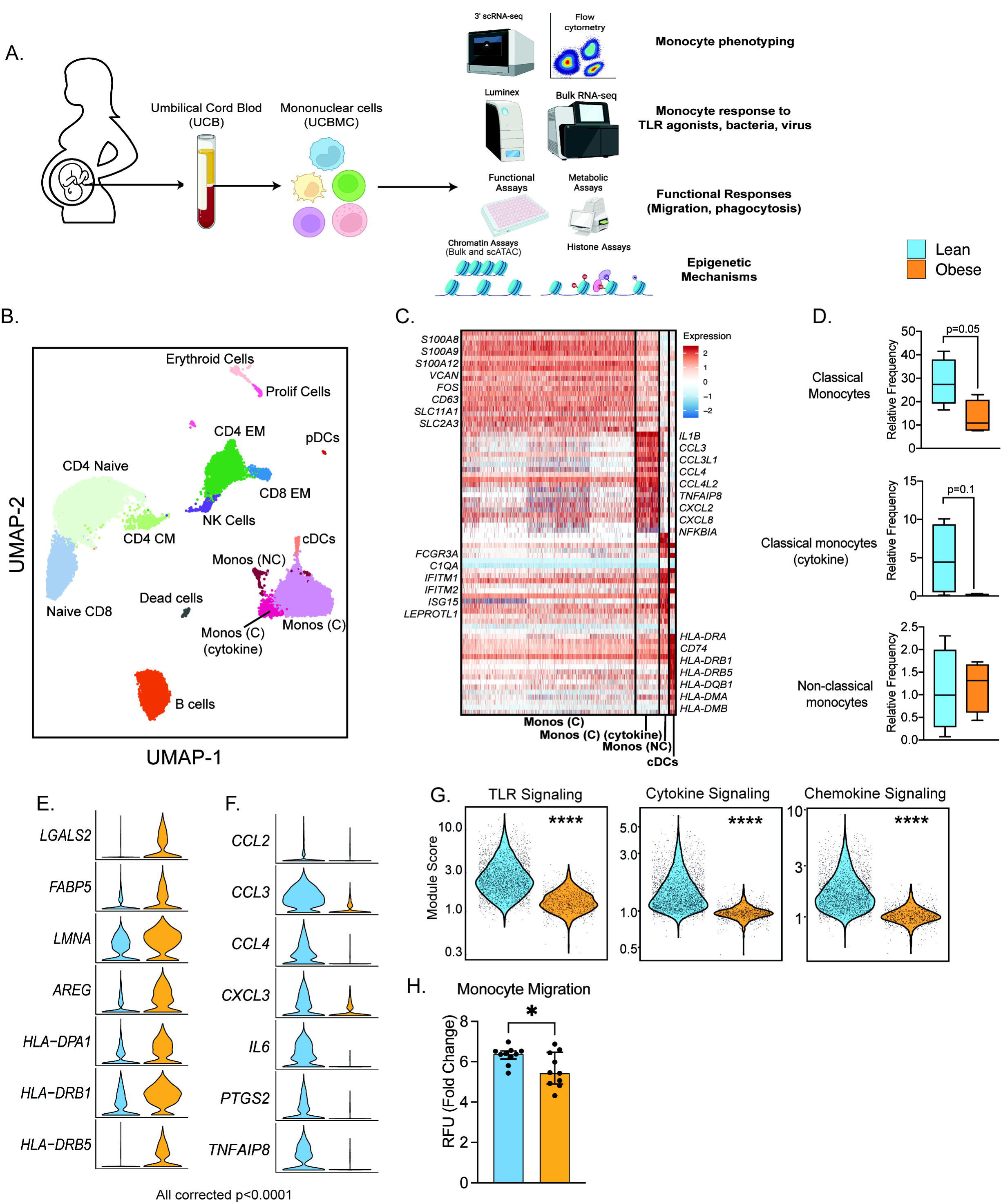
Experimental Design and phenotypic changes in UCB monocytes. (A) Experimental design for the study. Cord blood samples were obtained from neonates born to lean and mothers with obesity (n=43 lean and n=36 obese). All infants were delivered at term. Cohort characteristics are detailed in Table 1. UCBMC and plasma were isolated and used to assess the impact of pregravid obesity on fetal immunity using genomic and functional assays (flow cytometry, single-cell RNA and ATAC-Seq, ex vivo stimulations, phagocytosis, migration). (B) UMAP of term cord blood mononuclear cells collected from lean mothers and mothers with obesity (n=4/group). Samples were hashed using CellPlex (10X Genomics), sorted for live cells, and analyzed using a 10X single-cell 3’ gene expression assay. (C) Heatmap of top 30 markers of myeloid cell subsets (monocytes, conventional DCs) in UCBMC predicted using Seurat. (D) Box and whisker plots comparing relative frequencies of monocyte subsets (mean and ± SEM). (E-F) Violin plots of the top differentially expressed genes (FDR p<0.0001) in cord blood classical monocytes – I upregulated and (F) downregulated with maternal obesity. (G) Violin plots comparing module scores of TLR, cytokine, and chemokine signaling in cord blood classical monocytes with maternal obesity. (H) Bar graph comparing migration potential of UCB monocytes in response to supernatants from PMA-stimulated adult PBMC. Fold changes were calculated relative to no stimulation controls (mean and ± SEM).

Multi-parameter flow cytometry analysis of umbilical cord blood mononuclear cells (UCBMC) revealed a significant reduction in monocyte proportions (CD14+HLA-DR+) within live mononuclear cells (**Supp Figure 1D**). Additionally, within monocytes, pregravid obesity was associated with elevated non-classical monocyte subset (CD16+) (**Supp Figure 1D**). Despite this increase, UCB monocytes from the obese group responded poorly to ex *vivo* LPS (TLR4 agonist) stimulation (**Supp Figure 1D**), recapitulating the phenotype observed in previously reported studies (*15, 27, 28, 37*).

### Single-cell RNA sequencing reveals maternal obesity-associated transcriptional shifts in the UCB monocytes

We next asked if cord blood monocyte subsets were transcriptionally primed to respond poorly to *ex vivo* stimulation signals. To test this, we performed droplet-based single-cell RNA sequencing (scRNA-Seq) of UCBMC from lean and obese groups (n=4/group, hashed using Cell Plex). Principal Component Analysis (PCA) and Uniform Manifold Approximation and Projection (UMAP) analysis revealed the expected major immune subsets (**Figure 1B**) including T cells (*CD3D*, *CD8A*), comprised mostly of naïve cells (*CCR7*, *IL7R*) and some memory T cells lacking *CCR7*, B cells (*CD79A*, *MS4A1*), NK cells (*NCAM1*), conventional dendritic cells (cDCs, *HLA-DRA*, *FCER1A*), plasmacytoid dendritic cells (pDCs, *IRF8*), monocytes (*CD14*), erythroid cells (*HBB*), and a small subset of proliferating cells (*MKI67*) (**Supp Figure 2A).** A closer look at the monocytes revealed the presence of 3 clusters (all expressing CD14): classical monocytes (Mono C) expressing higher *S100A8*, *S100A9*, *VCAN*, non-classical monocytes (Mono NC) expressing higher *FCGR3A* and interferon-stimulated genes, and a cluster of classical monocytes (Mono (C) cytokine) with elevated levels of inflammatory transcripts *IL1B*, *CCL3*, *CXCL2*, and *CXCL8* (**Figure 1C**).

**Figure 2:**
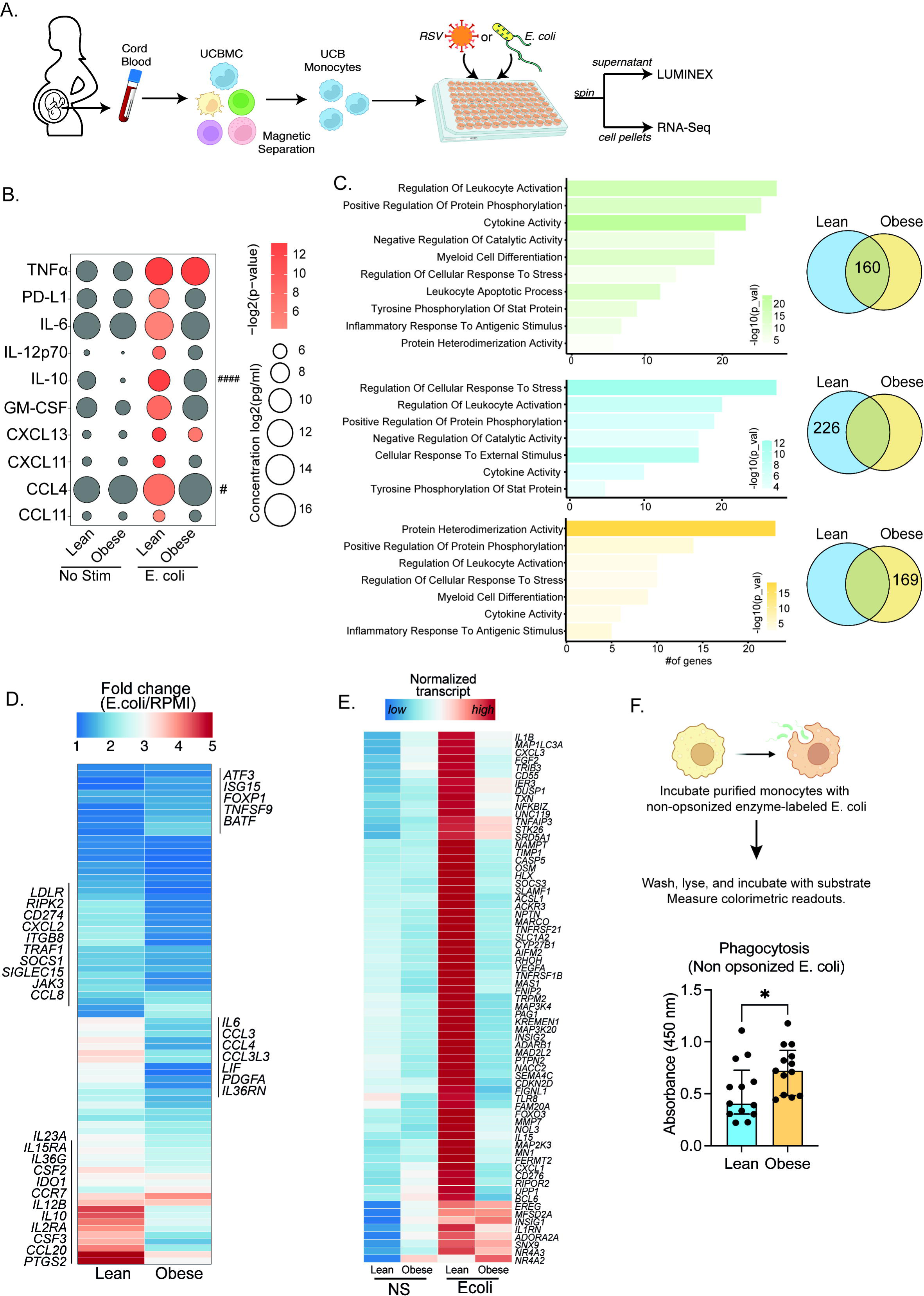
Cord blood monocyte responses to ex vivo *E. coli* infection are attenuated with maternal obesity) Experimental design for in vitro bacterial and viral stimulation. Purified monocytes were cultured in the presence/absence of *E. coli* or RSV for 16h. Cell pellets were used for bulk RNA-Seq analyses and supernatants were used for Luminex analyses of secreted cytokines and chemokines. (B) Bubble plot of key secreted factors significantly different following *E. coli* infection. The size of the bubble represents the quantity of the secreted analyte (log-transformed) whereas color represents statistical significance relative to no stimulation controls. Statistically significant analytes between lean and obese groups are highlighted with ^#^ - p<0.05. (C) Venn diagram (right) and corresponding functional enrichment (left) of genes upregulated with E. coli infection in lean and obese groups (Green denotes common DEG, blue DEG unique to lean group, and yellow DEG unique to the obese group). Bar colors represent the functional enrichment of DEGs predicted using Metascape. The length of the bar indicates the number of genes in each gene ontology (GO) term. Color represents the statistical significance of each GO term. (D) Heatmap comparing fold changes of the genes upregulated in both groups (77 genes) that mapped to GO terms “myeloid cell differentiation”, “inflammatory response to antigenic stimulus”, “leukocyte apoptotic process”, “regulation of leukocyte activity”, ”cytokine activity”, and ”positive regulation of protein phosphorylation”. (E) Heatmap comparing normalized transcript counts (blue – low to red – high) of genes exclusively upregulated in the lean group following *E. coli* infection. (F) Bar graph depicting colorimetric readout of phagocytosed *E. coli* particles by UCB monocyte. * or /^#^- p<0.05, ^####^ - p<0.0001

Comparisons of the relative abundance of these subsets revealed a decrease in classical monocytes (p=0.05) and a non-significant decrease in cytokine expressing monocyte clusters (p=0.1) with pregravid obesity (**Figure 1D** and **Supp Figure 2B**). This shift in UCB monocyte cell states is further demonstrated by differential gene expression analysis (**Figures 1E and 1F**) and module score comparisons (**Figure 1G**) within total UCB monocyte clusters. Pregravid obesity resulted in elevated expression of genes important for antigen presentation and immune regulation (**Figure 1E**). On the other hand, immune signatures associated with cytokine (*IL6*), chemokine (*CCL2*, *CCL3*, *CXCL3*), and TLR signaling were attenuated in the obese group (**Figures 1F and 1G),** in line with reduced TLR responses reported here (**Supp Figures 1D**) and prior studies (*28*). Dampened chemokine signaling translated to a reduction in the migration capacity of purified monocytes (**Figure 1H**).

### Pregravid obesity compromises cord blood monocyte responses to bacteria

Given the increased incidence of bacterial and viral infections in babies born to mothers with obesity (*22–24*), we next interrogated if the reduced UCB monocyte response to LPS stimulation (*15, 27, 28, 37*) extended to anti-microbial response. To that end, monocytes were purified from UCBMC and cultured with *E. coli* for 16 hours at 37°C. Secreted cytokines and chemokines were profiled using Luminex, while the transcriptional response to infection was recorded using bulk RNA sequencing (**Figure 2A**). While comparable levels of TNFL were secreted by both groups in absence of stimulus, an infection-induced increase in the secretion of pro-inflammatory cytokines (IL-6, IL12p70), regulatory cytokines (IL-10, PD-L1), chemokines (CCL4, CCL11, CXCL11) and growth factors (GM-CSF) was only detected in the lean group after *E. coli* stimulation (**Figure 2B**).

Next, we examined differences in transcriptional responses to *E. Coli*. The number of differentially expressed genes (DEG) relative to no stimulation was higher in the lean group (**Supp Figure 3A**). While ∼ 30% of genes up-regulated with infection (160 genes) were shared between the lean and obese groups (**Figure 2C**), the fold change of several of these genes involved in myeloid cell activation (*CSF2*, *CSF3*, *CCR7*) and cytokine signaling (*IL6*, *IL23A*, *IL10*, *IL12B*, *CCL2*, *CCL3*, *CCL4*, CCL8, *CCL20*) was lower in the obese group (**Figure 2D**). Importantly, functional enrichment of genes upregulated in the lean group alone revealed an over-representation of pathways associated with immune regulation (positive regulation of protein phosphorylation and regulation of cellular response to stress) (**Figure 2C**). This list included cytokines (*IL1B*, *IL1RN*), chemokines (*CXCL1*, *CXCL3*), growth factors (*VEGFA*, *FGF2*), and signaling molecules (*CASP5*, *NFKBIZ*, *SOCS3*, *TLR8*, *MMP9*) indicating a robust innate immune response in the lean group alone (**Figures 2E**). We also observed ∼50% overlap in genes downregulated with infection (**Supp Figure 3B**). These gene signatures enriched to GO terms associated with myeloid cell activation and negative regulation of differentiation (**Supp Figure 3C**). Interestingly, downregulation of MHC class II molecules (*HLA*-*DPA1*, *DPB1*, *DQB1*, *DRA*, *DRB1*, *DRB5*) was observed exclusively in the lean group (**Supp Figures 3C and 3D**).

**Figure 3:**
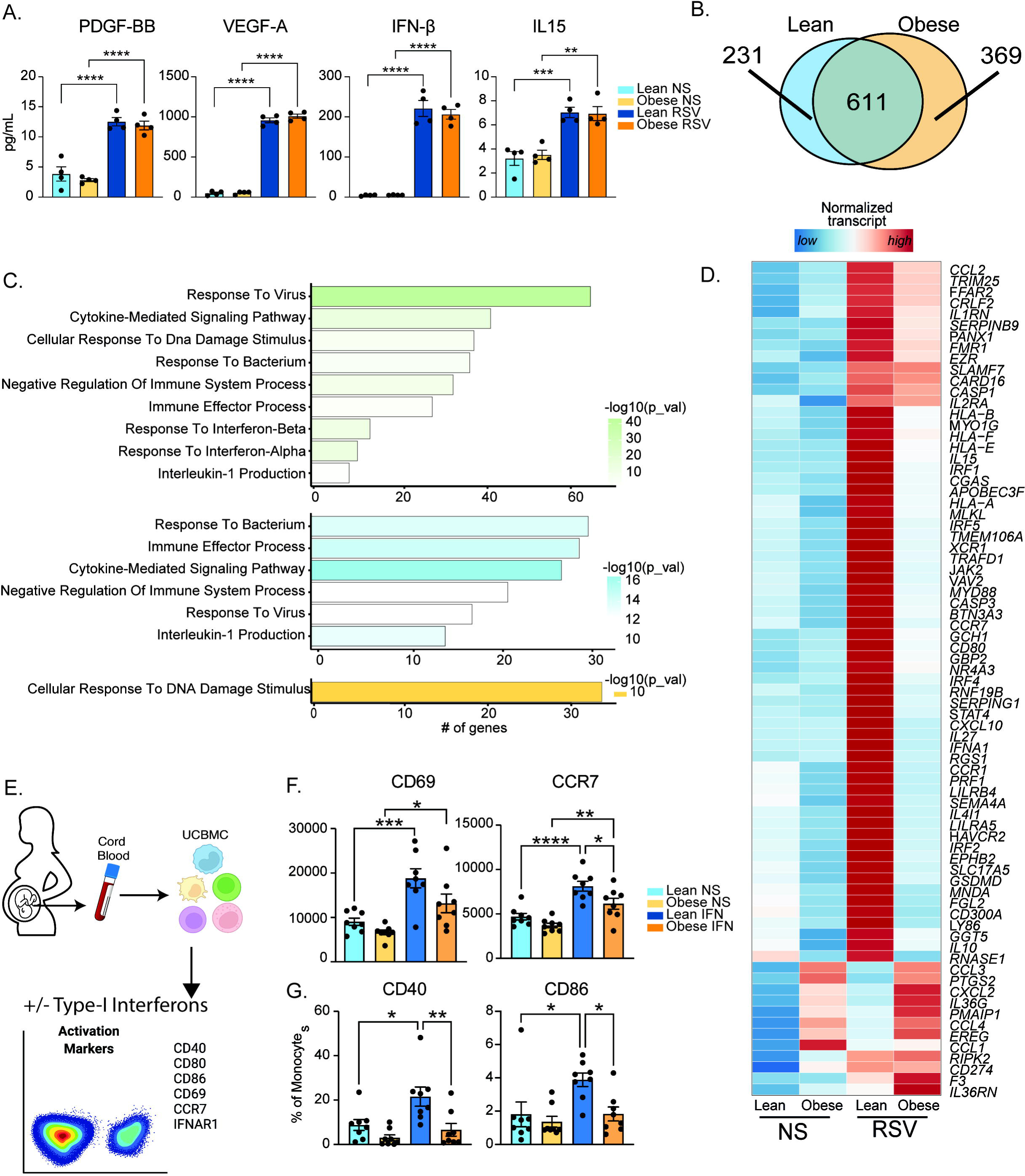
Maternal obesity does not alter acute anti-viral responses by cord blood monocytes but dampens responses to type-I interferon. A) Bar graphs comparing levels (mean and ± SEM) of key secreted factors by purified UCB monocytes in response to RSV infection measured using Luminex (n=4/group) compared to no stimulation (NS). (B) Venn diagram comparing DEG upregulated in response to RSV infection relative to no stim controls in lean and obese groups. (C) Functional enrichment of differentially expressed genes (DEG) detected in both groups (green, top), lean only (blue, middle), and obese group only (yellow, bottom) predicted using Metascape. (D) Heatmap comparing normalized transcript counts (blue – low to red – high) of genes exclusively upregulated in the lean group following RSV infection, mapping to GO terms “antigen processing and presentation”, “negative regulation of cell differentiation”, and “myeloid cell differentiation”. (E) Experimental design for testing monocyte responses to type-I IFN (n=8/group). UCBMC were stimulated with a mix of IFNα and IFNβ for 16 hours and activation markers were measured using flow cytometry (n=7-8/group). (F-G) Bar graphs comparing activation markers (F) CD69, CCR7, and (G) CD40 and CD86 in cord blood monocytes in response to type-I IFN. *- p<0.05, **- p<0.01, ***- p<0.001, ****-p<0.0001

Given the modest inflammatory response to *E. Coli* infection by UCB monocytes from the obese group, we next compared their phagocytic ability (**Figure 2F**). Purified UCB monocytes were cultured with labeled *E. coli* and probed for internalization. UCB monocytes from the obese group were more phagocytic compared to the lean group (**Figure 2F**), suggesting a more regulatory phenotype.

### Attenuated anti-viral transcriptional responses in UCB monocytes with pregravid obesity

Previous studies in rodent models have reported dysregulated inflammatory responses in the lungs of pups born to obese dams following RSV infection (*22*). Given that RSV is sensed by myeloid cells via TLR4/TLR8 pathways, we asked if maternal obesity compromises fetal myeloid cell responses to *ex vivo* RSV infection (**Figures 2A and 3**). Interestingly, we saw no differences between the lean and obese groups in terms of immune mediator production 16 hours post-stimulation (**Figure 3A**). UCB monocytes from the obese group generated a larger transcriptional response than those from the lean group (**Supp Figure 4A**). While gene signatures associated with a robust response to the virus were observed in both groups (**Figures 3B** and **3C**), significant differences in DEGs upregulated with RSV stimulation were noted (**Figures 3B**). Particularly, DEGs unique to the lean group were associated with anti-viral effector responses (**Figures 3C**). This list included genes involved in TLR signaling (*MYD88*, *GSDMD*, *CASP1*), cytokine and chemokine signaling (*CCL2*, *IL1RN*, *IL10*), and effector molecules that initiate a robust Th1 response (*STAT4*, *CXCL10*, *IL15*, *IL27*) to the virus (**Figure 3C and 3D**).

**Figure 4:**
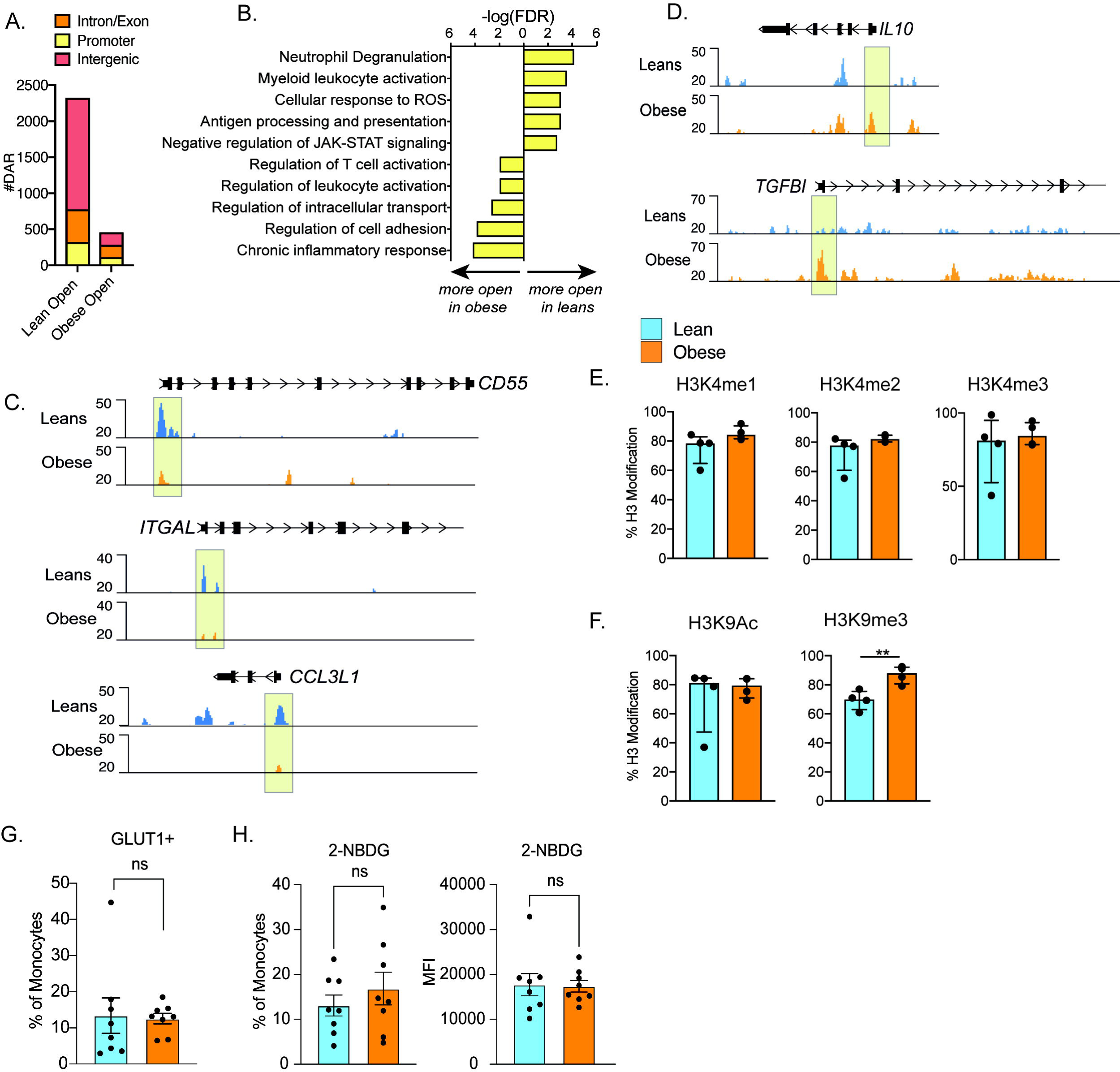
Epigenetic priming of cord blood monocytes with maternal obesity. (A) Stacked bar graphs comparing genomic contexts of differentially accessible regions (DARs) more open in either group. (B) Functional enrichment of genes regulated by DARs overlapping promoters using Metascape. (C-D) Representative pileups of genes more accessible in (C) lean or (D) obese groups. (E-F) Bar graphs comparing key histone modifications in (E) Histone H3 lysine 4 residue and (F) Histone H3 lysine 9 residues from nuclear extracts of resting UCB monocytes (n=3-4/group). Y-axis represents the percentage signal relative to the total H3 signal detected. (G) Comparison of surface glucose transporter GLUT1 and (H) glucose uptake using glucose analog 2-NBDG (n=7-8/group). Error bars represent medians with interquartile ranges. (** - p<0.01)

Interestingly, following RSV infection, several interferon-associated genes (ISGs) (*STAT4*, *IFNA1*, *IRF1*, and *IRF5)* were upregulated exclusively in the lean group (**Figure 3D**) while other ISGs were poorly induced in the obese group (**Supp Figure 4B**). Given comparable levels of secreted type-I IFN following 16h RSV infection (**Figure 3A**), these data suggested potential defects in the response to type I IFN in UCB monocytes from the obese group. To test this hypothesis, we stimulated total UCBMC with a cocktail of IFNL and IFNβ for 6 hours and measured activation markers using flow cytometry (**Figure 3E**). The frequency of cells expressing IFN receptor (IFNAR1) was comparable between the groups and treatment conditions as was the mean fluorescence intensity (MFI) following stimulation (**Supp Figure 4C**). Nevertheless, activation markers CCR7, CD40, and CD86 were upregulated to a lesser extent in the obese group (**Figure 3F**). Finally, analysis of DEG downregulated after RSV stimulation (**Supp Figure 4D**) in the lean group showed enrichment to transcription and translation while DEG unique to the obese group enriched to fatty acid and immunoglobulin binding (**Supp Figure 4E**).

### UCB monocytes from babies born to mothers with obesity are epigenetically and metabolically poised to respond poorly to pathogens

Prior studies from our laboratory have shown a dramatic loss of promoter methylation of several negative regulators of monocyte activation such as PPARL (*28, 38*). Given the prominent role of chromatin accessibility changes during acute responses to pathogens, we asked if UCB monocytes from babies born to mothers with obesity are epigenetically poised for dysfunctional responses to TLR ligands and pathogens. To test this, we isolated nuclei from purified resting CD14+ monocytes and identified global differences in baseline chromatin profiles using bulk ATAC-Seq. The analysis revealed significantly less open chromatin within promoter regions in the obese group relative to the lean group (**Figure 4A**). Genes regulated by promoters that were less accessible in the obese group were primarily involved in “myeloid cell activation”, “antigen processing and presentation”, and “neutrophil degranulation” (**Figure 4B**), which included proinflammatory *CD55*, *ITGAL*, and *CCL3L* (**Figure 4C**). In contrast, the few promoters that were more open in the obese group overlapped genes with regulatory roles such as “regulation of leukocyte activation” and “chronic inflammatory response” (**Figure 4B**), which included *IL10* and *TFGBI* loci (**Figure 4D**).

We next measured baseline differences in histone modifications that could potentially explain the differences in chromatin accessibility. Nuclear extracts from purified UCB monocytes were probed for specific methylation and acetylation signatures on histone H3K4 and H3K9 marks using ELISA. No differences in promoter-associated H3K4 mono-, di-, or tri-methylation were observed (**Figure 4E**). While H3K9 acetylation levels were comparable between the groups, heterochromatin associated H3K9 trimethylation was significantly elevated in UCB monocytes from the obese group (**Figure 4F**).

Finally, given these epigenetic differences at baseline, we interrogated the impact of pregravid obesity on the baseline energy demands of fetal monocytes. No differences were observed in surface expression of primary glucose transporter GLUT1 (**Figure 4G**) or overall uptake of extracellular glucose (**Figure 4H**).

### Epigenetic constraints to UCB monocyte activation with maternal obesity

We next asked if signaling defects downstream of TLR activation contribute to functional differences in UCB monocyte responses with maternal obesity. We began by measuring LPS- induced early phosphorylation events using flow cytometry. No differences in induction of MAPK-p38 or loss of IKBa phosphorylation signals were observed (**Figure 5A**). However, the increase in phosphorylation levels of NF-kB subunit p65 post LPS stimulation was less significant in the obese group (**Figure 5A**). To probe the impact of this modest difference, we measured nuclear NF-kB translocation using imaging flow cytometry. Fewer cells with translocated p50 were detected 4 hours following LPS stimulation (**Figure 5B**).

**Figure 5:**
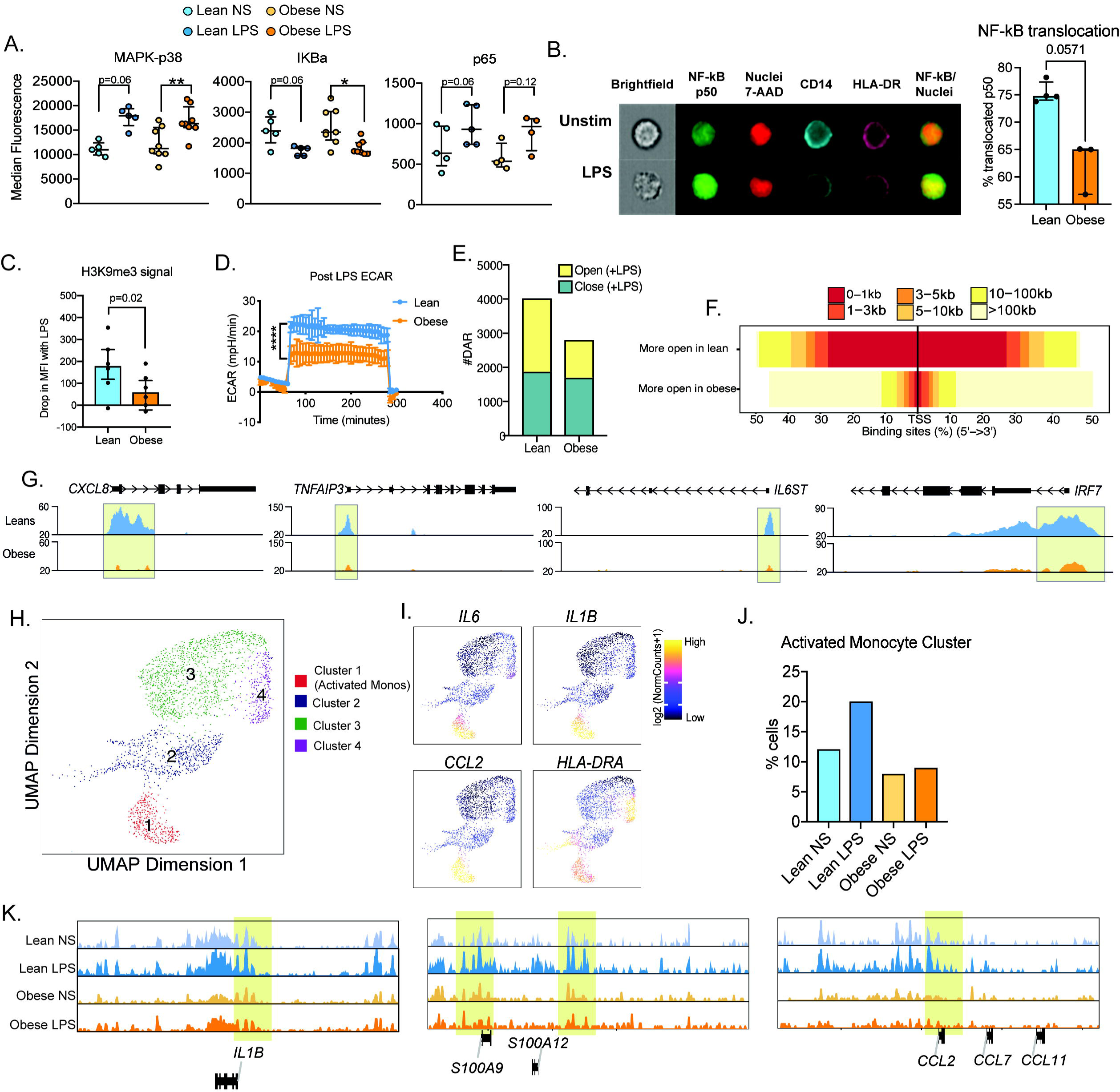
Epigenetic constraints to stimulation in cord blood monocytes from babies born to mothers with obesity. (A) Dot plots comparing median fluorescence intensity (MFI) ± SEM of phosphorylated signaling molecules downstream of TLR4 sensing (n=5/group). (B) Representative brightfield and fluorescent images of stimulated and unstimulated UCB monocytes profiled using imagine flow cytometry (n=3-4/group). NF-kB (p50-AF488) and nucleus (7-AAD) are shown in green and red respectively. Surface stains for CD14 and HLA-DR are shown in aqua and fuchsia respectively. Overlay of NF-kB and nuclei stain was used to determine translocation within CD14+HLA-DR+ monocytes. Bar graph comparing percentage translocated cells following LPS stimulation in lean and obese groups. (C) Bar graph comparing changes in trimethyl modification on H3K9 residues following LPS stimulation (relative to no stimulation controls) detected using flow cytometry. (D) Graph representing the kinetics of ECAR of stimulated monocytes following glucose injection and blockade of glycolysis. (E) Bar graph comparing DAR frequencies in each group following LPS stimulation. (F) Heatmap demonstrating overall accessibility differences following LPS stimulation around the promoter. (G) Pileups of key inflammatory loci post LPS stimulation (H) UMAP of single nuclei ATAC-Seq of LPS stimulated and sorted monocytes. (I) Feature plots demonstrating a cluster of activated monocytes (cluster 1). Color intensity represents fragments mapping to open chromatin regions. (J) Proportions of monocytes within each group mapping to activated monocyte cluster. (K) Pileups of inflammatory loci in activated monocytes with/without LPS stimulation. * - p<0.05; ** - p<0.01

We next asked if maternal obesity was associated with a dampened epigenetic remodeling in response to immune activation. Loss of the repressive of histone H3K9 trimethylation following LPS stimulation was reduced in UCB monocytes from the obese group (**Figure 5C**). Since metabolic changes precede these epigenetic changes, we next measured the overall increase in ECAR (a readout for glycolysis) following 1h LPS stimulation using a glycolytic stress assay. Following glucose injection, levels of ECAR increased significantly less in the obese group compared to the lean group (**Figure 5D**).

We next asked if suboptimal changes in chromatin accessibility correlate with attenuated responses to TLR activation in UCB monocytes. First, we compared chromatin accessibility via bulk ATAC-Seq on purified UCB monocytes following LPS stimulation. As observed in the resting state, fewer open regions were detected in the obese group following LPS stimulation (**Figure 5E**), primarily surrounding the transcription start sites (**Figure 5F**) and promoters of LPS-induced inflammatory molecules such as *CXCL8, IL6ST, TNFAIP3*, and transcription factor *IRF7* (**Figure 5G**).

To gain a better understanding of dampened epigenetic adaptations, we used single-cell ATAC-Seq of monocytes purified from resting or LPS stimulated (4 hours) UCBMC (**Supp Figure 5A**). We identified 4 major clusters, with cluster 1 clearly composed of activated monocytes (**Figure 5H**) where promoter regions regulating *IL6*, *IL1B*, *CCL2*, and *HLA-DRA* were significantly more open (**Figure 5I**). Overall, we observed a two-fold increase in the frequencies of activated monocytes in the lean group but no change in the obese group (**Figure 5J**). Moreover, as seen with the bulk ATAC-seq, overall changes in chromatin accessibility profiles within all monocyte clusters were more robust in the lean group (**Supp Figure 5B**). This included promoter regions of early inflammatory cytokine loci *IL1B*, chemokine *CCL2,* and alarmins *S100A9* and *S100A12* (**Figure 5K**).

### Poor fetal monocyte responses are recapitulated in a non-human primate model of western style diet (WSD)-induced maternal obesity

It is unclear if maternal obesity exerts the same impact on fetal monocytes cells *in utero* and if this hyporesponsive phenotype extends to tissue-resident macrophages. To address this, we leveraged access to fetal PBMCs, splenic, and gut (ileal) macrophages from rhesus macaques obtained at gestational day (GD) 130 from lean dams fed a control chow (CHOW) or obese dams fed a western diet (WSD) (**Figure 6A**). Despite a lack of difference in the frequency of circulating monocytes, the frequency of TNFL+IL-6+ producing monocytes obtained from the WSD group was significantly reduced following LPS stimulation (**Figure 6B**). Moreover, cytokine responses by splenic and ileal fetal macrophages stimulated with *E. coli* for 16hrs were also significantly dampened (**Figures 6C-F**). Principal component analysis shows that stimulation induced robust responses in fetal splenic (**Figures 6C**) and ileal macrophages (**Figures 6D**) from CHOW group. In contrast, stimulated macrophages from WSD group either overlapped or were very close to the non-stimulated condition (**Figures 6C and 6D**) along PC1. Specifically, following LPS stimulation, macrophages from the CHOW group produced significant increases in the levels of canonical Th1 polarizing factors (TNFL, IL-12, type I IFN, GM-CSF), Th17 factors (CCL20), and innate immune cell recruiting factors (CXCL2, IL-8, CXCL11) (**Figure 6E and 6F**). On the other hand, secretion of these factors was more modest in the WSD group following *E. Coli* stimulation. Consequently, levels of several immune mediators (TNFL, IFNL, IL-8, IL-10, CCL-20) were significantly lower than those observed in the CHOW group. These data highlight that maternal pregravid obesity leads to a significant dampening of fetal monocyte and tissue-resident macrophage responses to LPS and bacterial pathogens in primates.

**Figure 6:**
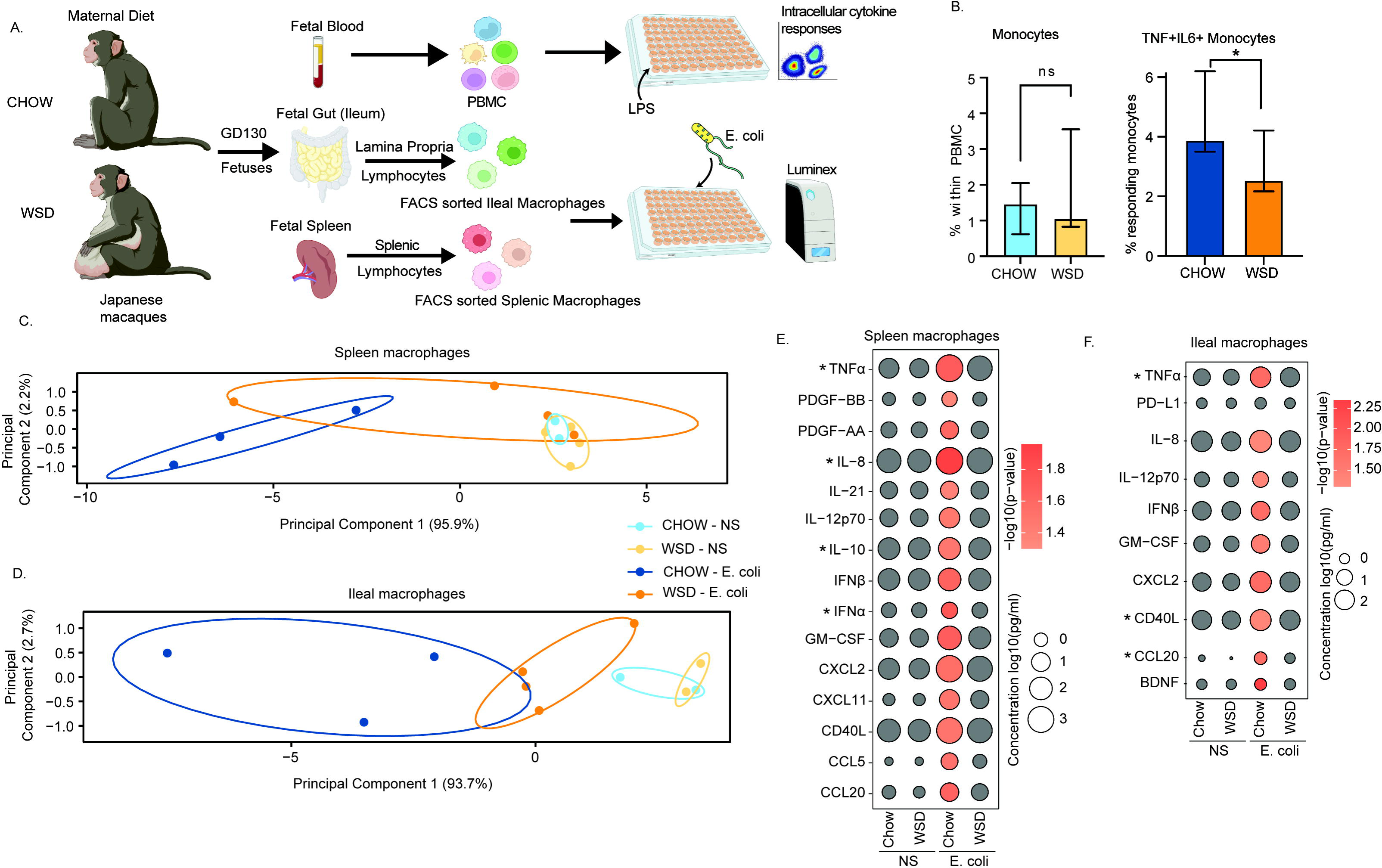
Maternal WSD attenuates fetal monocyte and macrophage cytokine responses to *ex vivo* stimulation. (A) NHP model of maternal diet-induced obesity. GD130 fetal gut (ileum) and spleen from rhesus macaque dams on a control chow diet (n=3) and western-style diet (n=4) were isolated. Fetal macrophages from ileal lamina propria lymphocytes and splenocytes were FACS-sorted and cultured overnight with *E. coli*. Supernatants were collected and cytokines/chemokines were measured using Luminex. (B) Bar graph representing the frequency (mean and ± SEM) of responding fetal monocytes to LPS in PBMC isolated from GD130 fetal circulation measured by intracellular cytokine staining. (C-D) Principal component analysis (PCA) of key secreted factors by unstimulated and LPS stimulated (C) splenic and (D) gut macrophages from chow and WSD groups. Each dot on the PCA corresponds to an aggregate (dimension reduced) of all cytokines/chemokines upregulated with stimulation. (E-F) Bubble plots comparing significantly different cytokine/chemokine levels in response to *E. Coli* stimulation in (E) splenic and (F) gut macrophages. Statistical differences between chow and HFD groups are highlighted. *-p<0.05

## DISCUSSION

Previous studies reported dampened responses by human UCB monocytes of babies born to mothers with obesity (*15, 28, 39, 40*) as well as by splenocytes from pups born to obese dams to LPS stimulation (*41*). Additional studies indicated increased airway reactivity and aberrant responses to RSV in pups born to dams fed a HFD during gestation (*22*). These observations align with clinical reports of increased admissions to neonatal intensive care units due to increased incidence of bacterial sepsis and enterocolitis (*17, 42, 43*). Mechanisms underlying this dysregulated response are emerging and indicate epigenetic rewiring in line with the Developmental Origin of Health and Disease (DOHaD) theory (*38, 44–50*). We confirm and extend these earlier observations in this study using an independent cohort of cord blood samples. Specifically, we investigated the impact of pregravid obesity on the epigenetic landscape and functional UCB monocyte response to bacterial and viral pathogens. Single-cell transcriptional analysis indicates a lack of a classical monocyte subset that contains high levels of cytokine/chemokine transcripts and is therefore poised to respond to stimulation. We further report a dysregulated response by UCB monocytes to viral and bacterial pathogens that are mediated by a remodeling of the chromatin landscape that leads to reduced accessibility at key inflammatory loci rather than defected in early TLR signaling events.

Data presented herein show that maternal obesity is associated with reduced classical UCB monocytes and more specifically, a subset expressing high levels of cytokine and chemokine transcripts (*IL1B, CCL3, CCL4, CXCL8, IL6*). Indeed, module scores for TLR/cytokine/chemokine signaling were all significantly reduced within UCB classical monocytes from babies born to mothers with obesity. These data are congruent with the dampened response to LPS observed previously (*15, 38*) and reduced migratory capacity of UCB monocytes from babies born to mothers with obesity. These observations differ from earlier bulk RNA-seq data (*28*) and highlight the power of single-cell transcriptomics to identify subtle defects at a much higher resolution. Importantly, these data indicate that UCB monocytes from babies born to mothers with obesity are predisposed towards a refractory/immune tolerant state.

We also report that responses to *E. coli* are disrupted with maternal obesity. Specifically, the production of key immune mediators was significantly dampened, and the magnitude and characteristics of the transcriptional response were altered. Notable differences include reduced expression of key inflammatory genes (while expression of negative regulators was increased). These responses are in line with prior *in vivo* rodent studies that showed increased susceptibility to bacterial challenge in pups born to obese dams (*26*). In contrast, UCB monocytes from the obese group exhibited increased phagocytic ability. Collectively, these observations support that maternal obesity skews fetal monocytes towards a regulatory phenotype. Indeed, prior studies have reported decreased levels of mRNA for pro-inflammatory cytokines IL-1β, and IL-12 in UCB monocytes from obese group compared to lean group (*30*). However, following differentiation, obese UCB monocyte-derived macrophages displayed an unbalanced response to M1 and M2 polarizing stimuli (*30*). These defects provide a potential explanation for increased susceptibility to bacterial infections in neonates born to mothers with obesity (*51, 52*).

Similarly, UCB monocytes from obese group showed a dysregulated response to RSV. Epidemiological studies have reported that high maternal BMI was associated with increased risks of offspring wheezing, prolonged cough, and lower respiratory tract infection (*53*). Additionally, studies in rodent models of maternal obesity showed that HFD dams delivered pups prone to develop more severe disease after RSV infection (*22*). Our data show a poor induction of key genes such as interferon response factors, co-stimulation molecules, chemokine receptors, and numerous ISG. Intriguingly, poor induction of ISG occurred despite comparable protein levels of IFNβ. It is possible that lower transcript levels of *IFNA* could contribute to this outcome. However, other factors are likely to contribute given the dampened ability of the cells to respond to IFNL/IFNβ when supplied *in vitro*.

Collectively, these data suggested a disruption in the ability of UCB monocytes from the obese group to interpret signals received through PRR. Prior studies (*28*) and data enclosed in this study show that TLR and IFN receptor surface protein expression was comparable between the two groups. Moreover, activation of key signal transduction proteins downstream of TLR4 was also comparable between the two groups. However, nuclear translocation of the key transcriptional factor NF-kB was reduced with maternal obesity. While chromatin accessibility regulates NF-kB binding and subsequent induction of gene expression, NF-kB binding also influences the chromatin state by recruiting chromatin-modifying enzymes and evicting negative regulators (*54*). Indeed, our data suggest a significant reduction in chromatin accessibility in UCB from the obese group that was accompanied by an increased abundance of a repressive histone mark (H3K9me3) that persisted upon LPS stimulation. Single-cell ATAC-Seq revealed the presence of an activated monocyte cluster, the abundance of which increased in the lean group but not the obese group further supporting the inability to properly integrate signals following signaling through pattern recognition receptors. Closed chromatin profiles at baseline suggest that these cells are epigenetically poised to respond poorly to subsequent stimulation. Thus, the lack of robust metabolic, epigenetic, and transcriptional response to stimulation provides further evidence for mechanistic constraints contributing to innate immune tolerance.

Analysis of circulating monocytes and tissue-resident macrophages obtained from third- trimester rhesus macaque fetuses showed similar outcomes as those reported for human umbilical cord blood samples, with dampened responses to LPS and *E. coli.* These data strongly suggest that the defects we observed are not limited to UCB, but rather that maternal diet leads to in utero re-programming of the developing immune system. More importantly, the fact that both circulating monocytes and tissue-resident macrophages generated a dampened response strongly suggests that this programming occurs early during gestation. Indeed, monocytes and macrophages are some of the earliest immune cells to develop with the first wave originating from late yolk sac-derived erythromyeloid progenitors and fetal liver (*55*). Fetal macrophages of the gut are believed to be originally derived from fetal liver monocytes that are replaced later in life by bone marrow-derived monocytes (*55*). Circulating and splenic macrophages in late gestation are a mix of fetal liver and bone marrow-derived monocytes (*55*). Signals from the maternal microenvironment influence monocyte production and their functional programming. The heightened maternal inflammatory milieu during critical windows of fetal immune ontogeny may skew the maturation of immune cells towards a regulatory state, perhaps a protective adaptation to limit fetal inflammation.

The transgenerational impact of maternal obesity on the epigenome of the offspring’s immune system has been long appreciated. Several studies have highlighted alterations in methylome and chromatin accessibility landscape of the offspring’s circulating immune cells that are evident as early as at birth (*15, 27, 38, 48, 56*). Alterations in promoter regions of DNA methylation in fetal monocytes parallel their dampened expression of anti-inflammatory mediators and attenuated response to M1/M2 polarizing stimuli (*30*). We previously reported that maternal obesity is associated with baseline global hypomethylation at promoter regions of UCB monocytes (*28*). These differentially methylated regions were found to be involved in metabolism, cell migration and adhesion of myeloid cells, immune development, and defense/stress responses and overlapped with inactive promoters and open chromatin (*28*).

Mechanisms that underlie these baseline epigenetic changes are poorly understood. Some of these epigenetic modifications may be inherited through alterations in gametes, but others may be in response to maternal inflammatory and nutritional cues (*57*). For instance, the frequency of non-classical monocytes is increased in UCB from babies born to mothers with obesity, suggesting *in utero* activation. It is unclear how this activation may be happening since although pregravid obesity leads to significant differences in levels of maternal circulating inflammatory immune mediators, only leptin levels are significantly increased in UCB plasma. A neurotropic factor shown to be important for early cognitive and behavioral development, leptin resistance is associated with long-term metabolic and behavioral outcomes. Altered leptin signaling *in utero* may predispose the fetus to leptin resistance and possibly be the link to the high rates of fetal overgrowth and obese offspring born to mothers with obesity (*58*). Indeed, reports show a positive correlation between cord blood leptin concentrations, birth weight (*59*), and the risk of developing asthma later in childhood (*60*). Furthermore, leptin has been shown to stimulate monocyte activation and differentiation (*61*). These observations provide one plausible avenue for future mechanistic exploration of maternal-fetal communications that result in fetal immune cell reprogramming with maternal obesity. Nevertheless, the dysregulated fetal monocyte phenotype under ex vivo conditions highlights an evolutionary adaptation to protect the fetus from exacerbated inflammatory responses and subsequent tissue damage. However, this regulatory skewing also contributes to less-than-ideal responses against pathogens, as observed in both animal models and human studies.

### Data availability

The datasets supporting the conclusions of this article are available on NCBI’s Sequence Read Archive PRJNA847067. https://dataview.ncbi.nlm.nih.gov/object/PRJNA847067?reviewer=jaohha21p6li1t89aresdn73ms

### Competing interests

The authors have declared that no financial or non-financial competing interests exist.

## Funding

This study was supported by grants from the National Institutes of Health 1K23HD06952 (NEM), R03AI112808 (IM), 1R01AI142841 (IM), and 1R01AI145910 (IM).

## Author Contributions

SS, NEM, and IM conceived and designed the experiments. SS, NM, and BMD performed the experiments. SS, BMD, and NM analyzed the data. SS, NEM, and IM wrote the paper.

## Acknowledgments

We thank Mr. Allen Jankeel and Ms. Gouri Ajith for help with immune assays and the preparation of RNA-Seq libraries. We thank Dr. Jennifer Atwood from the flow cytometry core at the Institute for Immunology, UCI for assistance with sorting experiments and imaging flow cytometry. We thank Dr. Melanie Oakes from UCI Genomics and High-Throughput Facility for assistance with 10X library preparation and sequencing.

## SUPPLEMENT FIGURES

**Supplementary Figure 1: Longitudinal changes in circulating inflammatory environment.** (A) Complete blood cell counts of umbilical cord blood samples obtained from babies born to 34 lean or 26 mothers with obesity. (B) Circulating levels of CRP and IL-6 were measured using an ELISA. (C) Circulating levels of insulin and leptin were measured using a metabolic Luminex panel. (D) The frequency of monocytes and non-classical monocyte subsets as well as IL- 6+TNFa+ producing monocytes was determined using flow cytometry.

**Supplementary Figure 2: Umbilical cord blood mononuclear cell profiling using scRNA-Seq.** (A) Violin plot delineating markers used to define major immune cell subsets in cord blood samples. Candidate genes were selected from the list of top markers predicted by *FindAllMarkers* function in Seurat (B) UMAP of UCBMC profiles colored by the group.

**Supplementary Figure 3: Cord blood monocyte responses to *E. coli*** (A) Bar graph representing the number of upregulated (red) or downregulated (blue) differentially expressed genes (DEG) in each group. (B) Venn diagram representing the number of downregulated DEG following *E. coli* stimulation in each group. (C) Functional enrichment of downregulated DEG detected in both groups (top, green), or exclusively in the lean (middle, blue) and obese (bottom, orange) groups. (D) Violin plots comparing fold changes of DEGs downregulated following *E. coli* stimulation and involved in antigen presentation.

**Supplementary Figure 4: Cord blood monocyte responses to RSV.** (A) Bar graph representing the number of upregulated (red) or downregulated (blue) DEG in each group. (B) Heatmap of upregulated genes involved in interferon signaling pathway. (C) Surface protein levels of interferon receptor 1 (IFNAR1) represented as a fraction of cells expressed (top) or median fluorescent intensity (bottom) (D) Venn diagram representing the number of downregulated DEG following RSV stimulation (E) Functional enrichment of downregulated DEG detected in both groups (top, green), or exclusively in the lean group (top, blue) and obese group (bottom, orange).

**Supplementary Figure 5: Epigenetic responses to LPS in cord blood monocytes.** UMAP of single-cell chromatin profiles of sorted cord blood monocyte nuclei. Clusters represent the pool of nuclei from lean and obese groups before and after stimulation. (B) MA plots comparing fold changes in chromatin accessibility changes following LPS stimulation in lean (left) and obese (right) groups with upregulated genes in red, downregulated genes in blue, and not differentiated genes in grey.

## Notes

### Competing Interest Statement

The authors have declared no competing interest.

