## Supplementary figures and images for "Impact of pregravid obesity on anti-microbial fetal monocyte response"

### Supp Figure 1

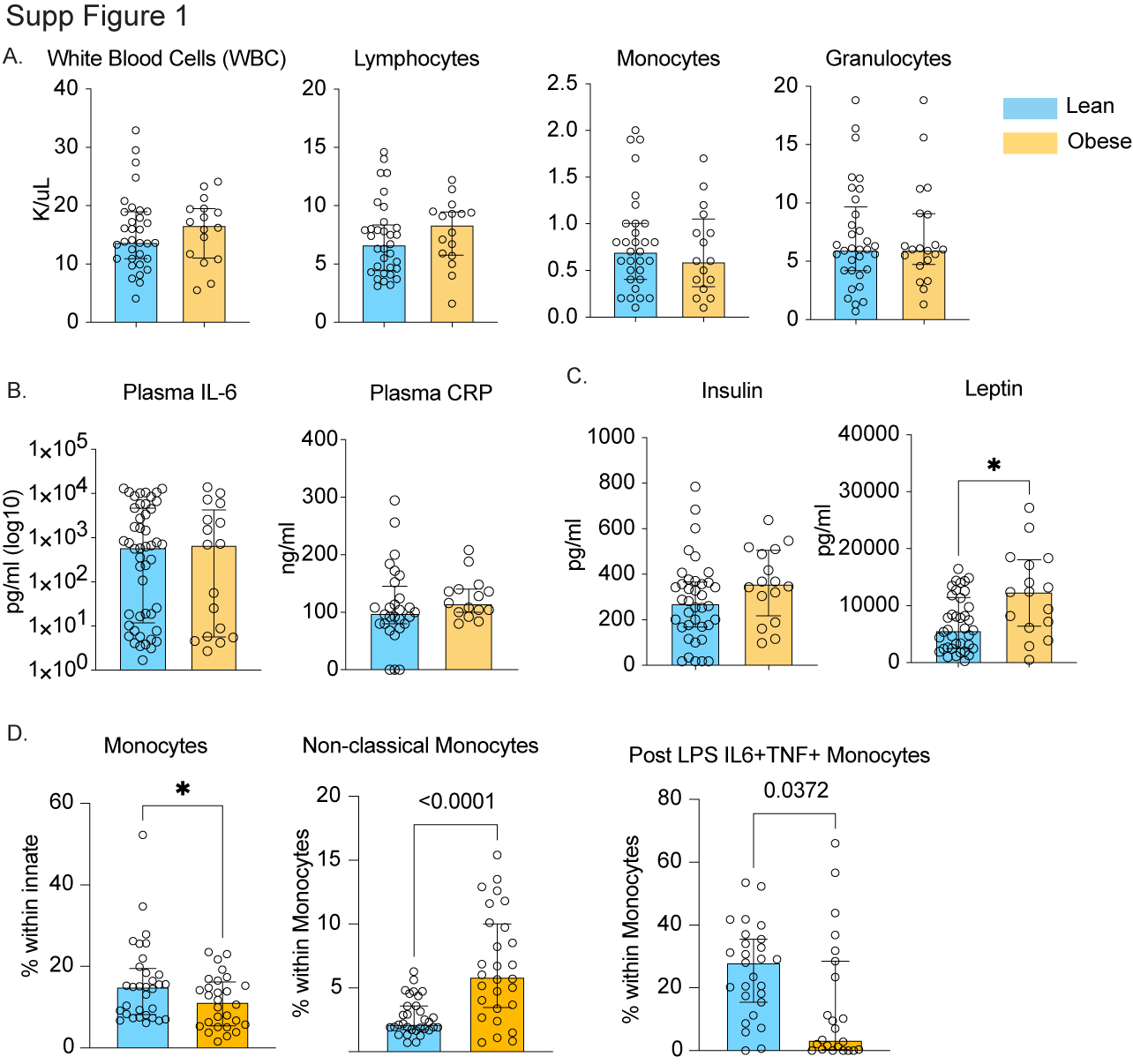

### Supp Figure 2

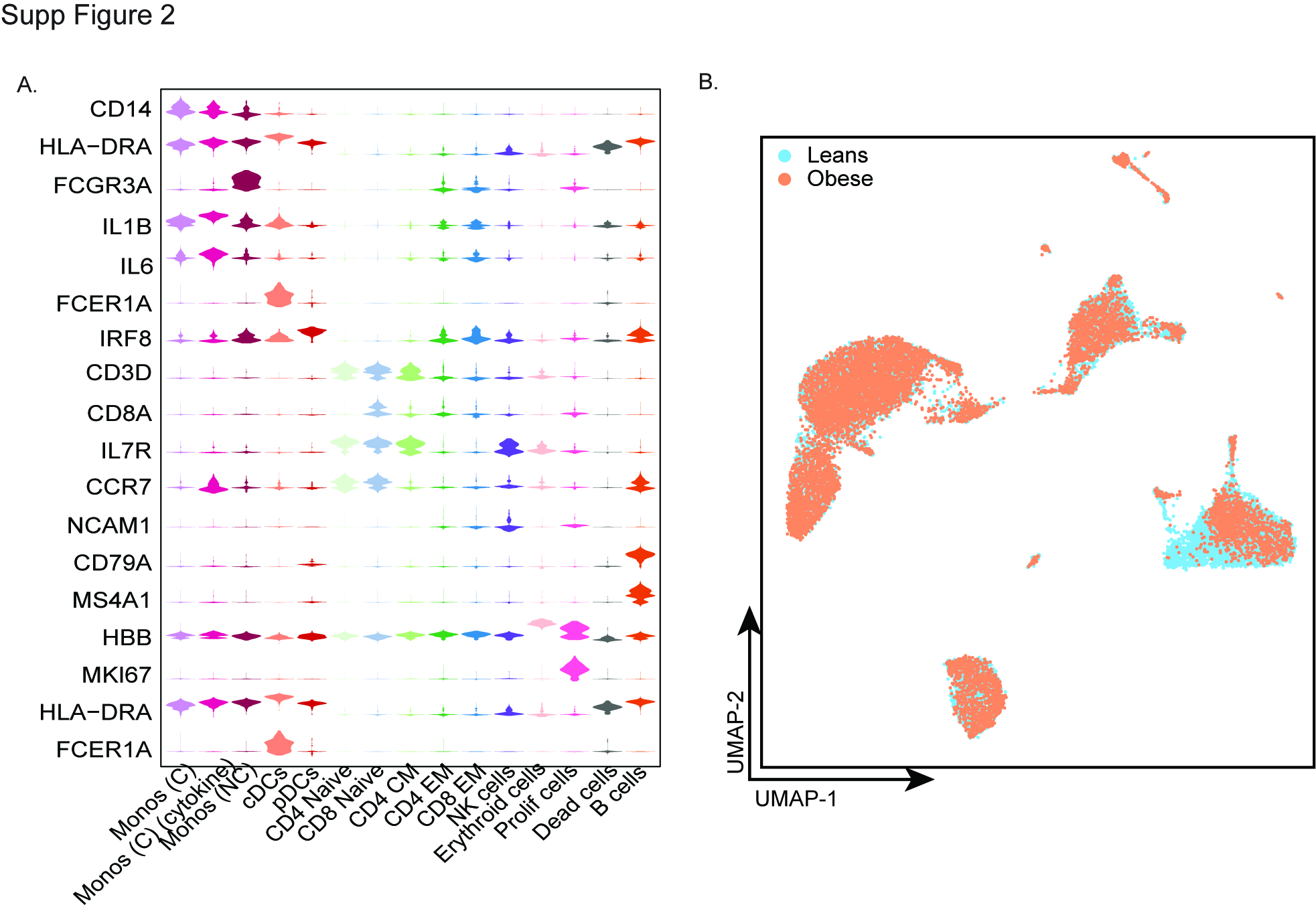

### Supp Figure 3

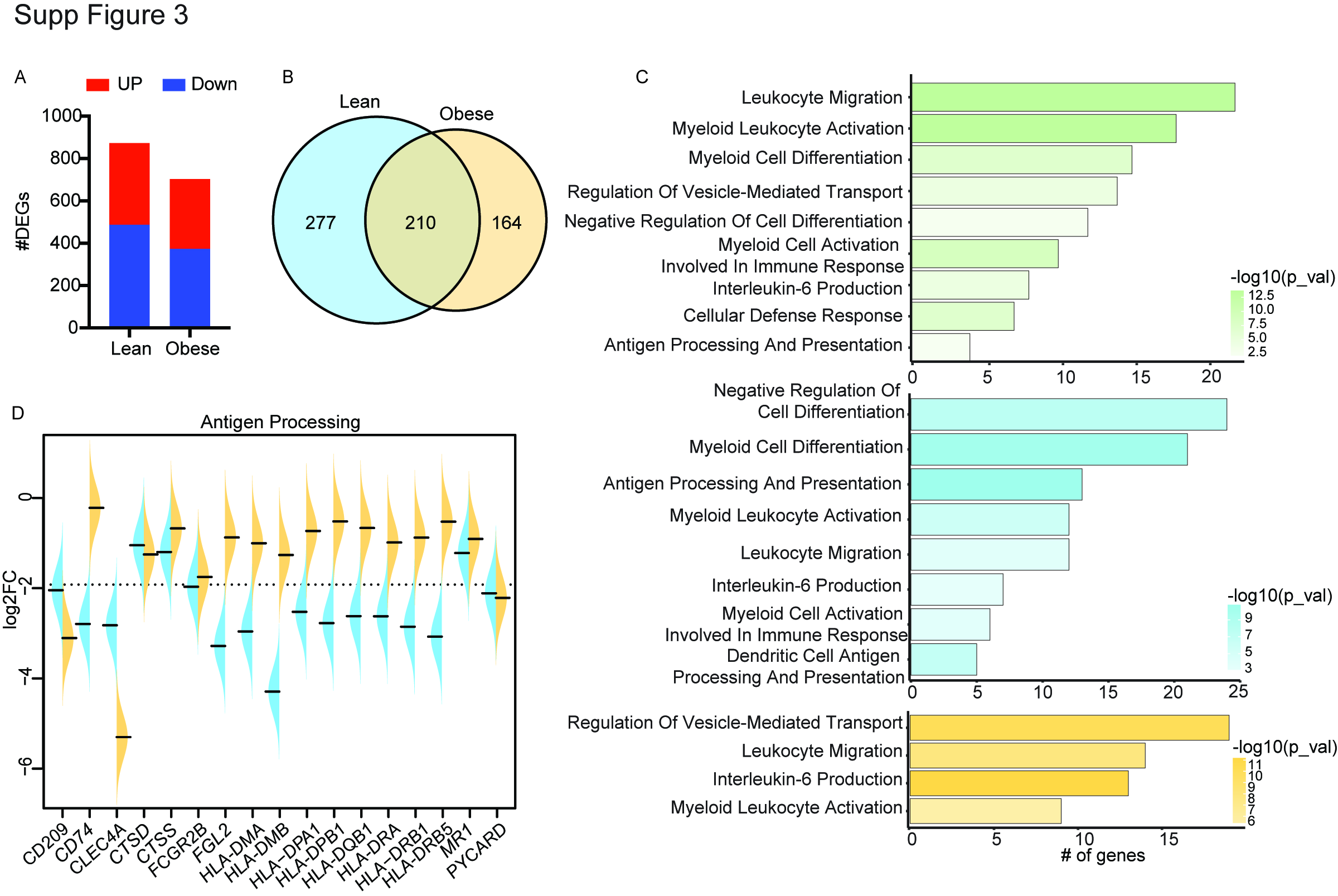

### Supp Figure 4

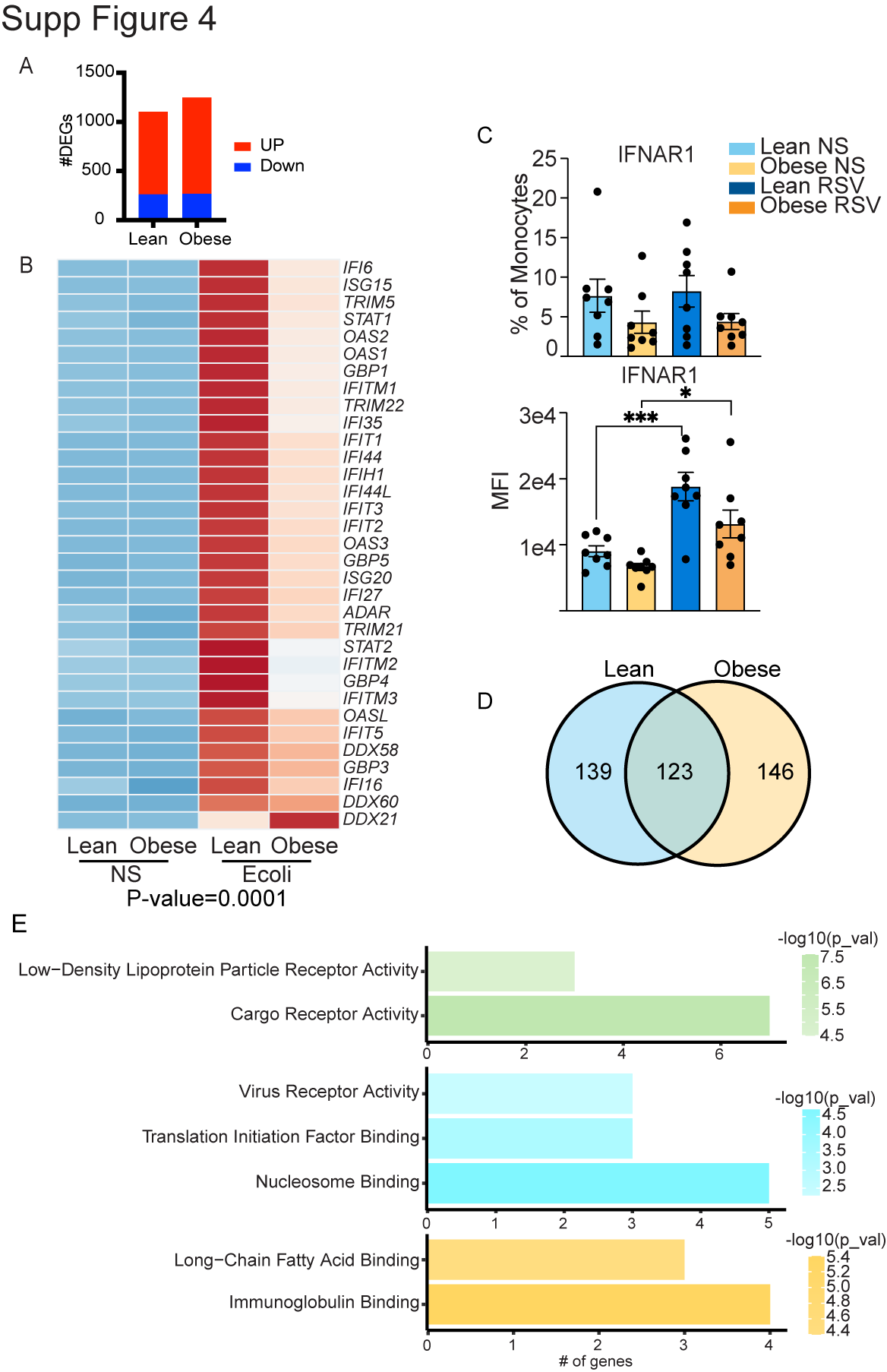

### Supp Figure 5

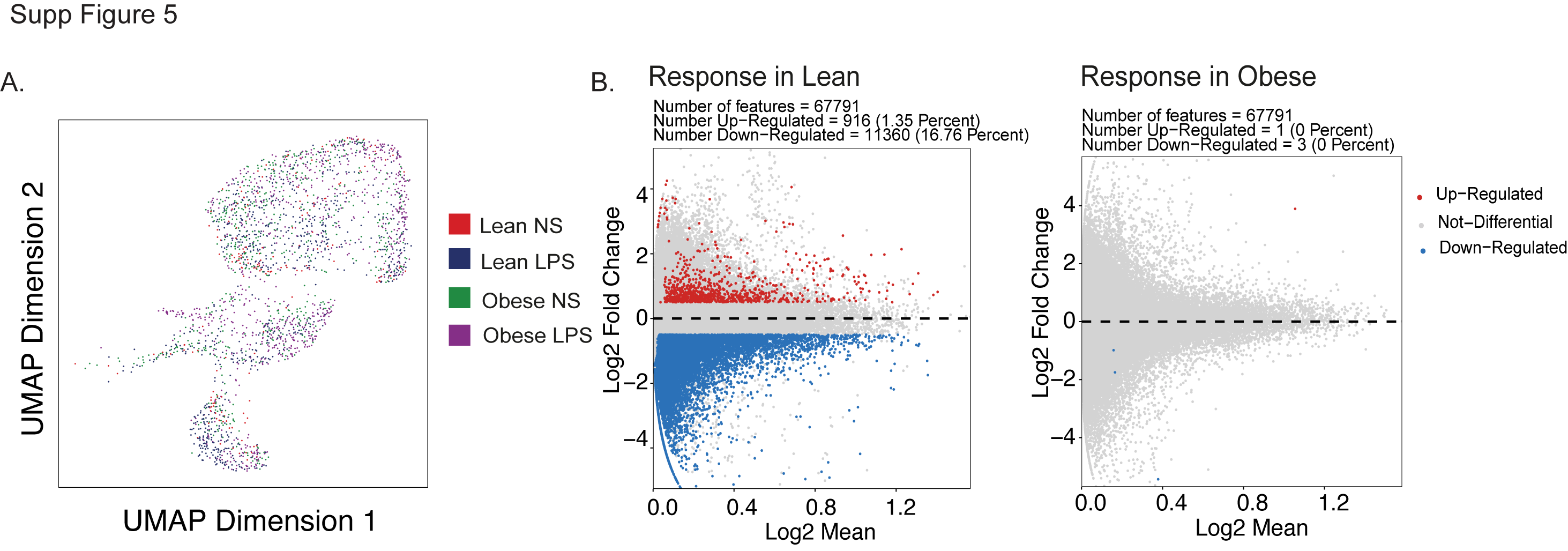
